# Temporal transcription factors determine circuit membership by permanently altering motor neuron-to-muscle synaptic partnerships

**DOI:** 10.1101/2020.03.25.007252

**Authors:** Julia L. Meng, Yupu Wang, Robert A. Carrillo, Ellie S. Heckscher

## Abstract

Previously, using the Drosophila motor system as a model, we found the classic temporal transcription factor, Hunchback acts in NB7-1 neuronal stem cells as a molecular switch to control which circuits are populated by NB7-1 neuronal progeny (Meng et al., 2019). Here, we manipulate cardinal transcription factors, Nkx6 and Hb9, which are candidate effectors of Hunchback and which alter axon pathfinding in embryos. Yet manipulation of these cardinal transcription factors does not permanently alter neuromuscular synaptic partnerships. This demonstrates that compensation can correct early defects. We perform additional temporal transcription factor manipulations, precociously expressing Pdm and Castor in NB7-1 and prolonging expression of Hunchback in NB3-1. In every case, we find permanent alterations in neuromuscular synaptic partnerships. These data support the idea that temporal transcription factors are uniquely potent determinants of circuit membership, which do not trigger compensatory programs because they act to establish the expected pattern of wiring for the motor system.

## Introduction

During neural development, neurons are born from neuronal stem cells. Neurons then make a series of decisions that culminate in the assembly of neuronal circuits. For example, motor neurons first decide to send axons out of the CNS; then axons decide to travel to a particular muscle field; once within a muscle field, axonal growth cones decide which muscle fiber to contact, and finally growth cones decide to form functional synapses. Circuit membership of neurons is determined by the series of decisions they make. How these decisions are regulated is a key question in developmental neurobiology.

The class of molecules termed temporal transcription factors are poised to be key upstream regulators of the decisions that determine circuit membership. This is because in many systems there is a strong association between the birth time of a neuron and its ultimate circuit membership, and because temporal transcription factors have been postulated to control entire time-linked developmental programs (Bhansali et al., 2014; Deguchi et al., 2011; Eerdunfu et al., 2017; Greaney et al., 2017; Jefferis et al., 2001; Kulkarni et al., 2016; McLean et al., 2007; McLean and Fetcho, 2009; Morrow et al., 2008; Osterhout et al., 2014; Petrovic and Hummel, 2008; Pujol-Martí et al., 2012; Tripodi et al., 201; Doe 2017, Li et al. 2013, Suzuki et al. 2013, EIliott et al., 2008, Alsio et al., 2013, Allan and Thor, 2015). If temporal transcription factors control entire time-linked developmental programs, this would include all of the decisions that a neuron makes during development to establish circuit membership. However, these decisions occur in the context of a dynamic system, the developing animal, and so, dynamic environmental factors have the potential to strongly influence circuit membership. In many brain regions and many model systems, temporal transcription factors control marker gene expression in newly-forming, post-mitotic neurons (Isshiki et al. 2001, Moris-Sanz et al. 2014, Novotny et al. 2002, Pearson & Doe 2003, Tran & Doe 2008, Cleary & Doe 2006, Grosskortenhaus et al. 2006, Baumgardt et al. 2009, Stratmann et al. 2016). Temporal transcription factors also control neurotransmitter expression in embryonic neurons (Stratmann and Thor, 2017; Isshiki et al., 2001, Allan and Thor, 2015). In Drosophila mushroom body, central complex, and nerve cord, temporal transcription factors control axon and dendrite targeting and neuron morphology (Pearson and Doe, 2003; Seroka and Doe 2019; Meng et al., 2019, Sullivan et al., 2019; Rossi et al., 2017; Doe et al., 2017). These data demonstrate temporal transcription factors regulate the early decisions a neuron makes towards circuit membership.

It is still poorly understood how manipulation of temporal transcription factors impacts final neuronal decisions--synaptic partner choice and synapse formation--which are the critical final steps of determining circuit membership. Notably, synapse formation is distinct from other aspects of neuronal maturation in that it is obligately non-cell autonomous because it requires two permissive partners cells in order to form a functional synapse. A recent study in *Drosophila* central complex showed that loss of the temporal transcription factor, Eyeless/Pax6, altered the number of neurons produced, and impacted the adult navigation behavior, demonstrating a long-lasting change in navigational circuits (Sullivan et al., 2019). Although Eyeless controls morphology, it is unknown how Eyeless impacts synaptic partner selection and function of altered neurons. Another recent study in Drosophila nerve cord showed that prolonged expression of the temporal transcription factor, Hunchback, in one neuronal stem cell, <u>n</u>euro<u>b</u>last (NB) 7-1 impacts functional synaptic connections made by NB7-1 progeny (Meng, et al. 2019). Specifically, expression of Hunchback increases the total number of Even-skipped expressing (herein: Eve(+)) motor neurons made by NB7-1 without significantly extending the total number of neurons produced in the NB7-1 lineage. In wild type larvae, Eve(+) motor neurons exit the CNS, travel to the dorsal muscle field and make functional synaptic connections on a subset of dorsal muscles (Meng et al., 2019), and the ectopic Eve(+) motor neurons generated by prolonged expression of Hunchback also project to and form synaptic connections with muscles in the dorsal muscle field (Meng et al., 2019). Therefore, Hunchback expression in neuroblast, NB7-1 determines how many of the lineage’s neuronal progeny populate the dorsal muscle circuit. These data unequivocally demonstrate that Hunchback activity in NB7-1 is a key regulator of motor circuit membership. However, it is still unknown to what extent regulation of circuit membership is a general feature of all temporal transcription factors, and to what extent this feature is widespread and extends to other classes of transcription factors. For example, it is unknown to what extent the temporal transcription factor Hunchback can control circuit membership of neuronal progeny from any neuroblast; to what extent temporal transcription factors other than Hunchback can control circuit membership; and to what extent other classes of transcription factors can control circuit membership.

In addition to temporal transcription factors, a second class of transcription factors called cardinal transcription factors are prime candidates to be regulators of circuit membership because these factors regulate motor axon outgrowth to specific muscle groups in embryonic stages (Landgraf et al., 1999; Broihier et al., 2002; Broihier et al., 2004; Labrador et al., 2005). Cardinal transcription factors are highly conserved, are found across many brain regions and model systems, and include factors such as EN1/engrailed, HB9, EVX1/Eve, SIM1/Sim (Lu et al., 2015; Arber 2012; Briscoe et al., 2000; Jessell 2000; Pierani et al., 2001; Gribble et al., 2007). Cardinal transcription factors are distinguished from temporal transcription factors in several ways. First, temporal transcription factors are thought to specify the temporal identity of a neuron (e.g., first-born, second-born), regardless of stem cell origin (e.g., NB7-1, NB3-1), and regardless of cell type (e.g., motor neuron, glia), whereas cardinal transcription factors are known to specify cell type (e.g., dorsal motor neuron, ventral motor neuron, commissural interneuron, glia, etc.). Second, temporal transcription factors act in neuronal stem cells, whereas cardinal transcription factors act in newly-formed, post-mitotic neurons (Hirono et al., 2017; Broihier et al., 2002; Broihier et al., 2004; Landgraf et al., 1999; Cheesman et al., 2004). Third, temporal transcription factors are thought to be nested in complex loops of activation and inhibition (Nakajima et al., 2010), whereas cardinal transcription factors have cross-repressive interactions (Briscoe and Ericson, 2001; Broihier et al., 2002; Broihier et al., 2004; Landgraf and Thor, 2006). Cardinal transcription factors have been shown to regulate axon guidance in maturing motor neurons (Landgraf et al., 1999; Broihier et al., 2002; Broihier et al., 2004, Cheesman et al., 2004). These data suggest that in motor neurons, cardinal class transcription factors regulate the early decisions important for determining circuit membership. An attractive hypothesis is that cardinal transcription factors are downstream effectors of temporal transcriptions acting as key regulators of circuit membership (Seroka et al., 2020 *Biorxiv*). To test this model, we manipulate expression of cardinal transcription factors and examine circuit membership (i.e., synaptic connectivity) in mature circuits.

In this study, we ask to what extent is regulation of circuit membership a feature of the class of temporal transcription factors. We manipulate temporal transcription factors and cardinal transcription factors and characterize the final steps of circuit membership--synaptic partner selection and functional synapse formation using the Drosophila larval motor system as a model. The Drosophila motor system is organized into anterior-posterior repeated, left-right symmetrical hemisegements (Figure 1A-E). Each hemisegment contains 30 muscle cells, which are organized into three main groups—dorsal, transverse, and ventral (Figure 1B). Each muscle group is controlled by a unique circuit (Bate, 1993; Heckscher et al., 2012; Zarin et al., 2019), and populating each circuit are non-overlapping sets of motor neurons. Importantly, by late larval stages Drosophila motor neuron-to-muscle synapses are large, experimentally-accessible, and extremely well-characterized (Nose, 2012) (Figure 1E). Using this system, we determine how temporal transcription factors and cardinal transcription factors impact circuit membership by examining motor neuron-to-muscle synaptic partnerships. First, we find that loss of the cardinal transcription factors, nkx6 and hb9, does not change motor neuron-to-muscle synaptic partnerships at late larval stages, despite the fact that during embryonic development loss of nkx6 and hb9 dramatically alters axon guidance (Broihier et al., 2004; Cheesman et al., 2004). These data suggest the existence of compensation that can repair early developmental defects. These findings are in contrast to our previous work showing that the temporal transcription factor, Hunchback acting in NB7-1 can permanently alter motor neuron-to-muscle synaptic partnerships (Meng et al., 2019). This contrast highlights Hunchback’s potency to regulate circuit membership. Second, we find that Hunchback acting in NB3-1 is able to specify synaptic partnerships and therefore circuit membership of NB3-1 neuronal progeny. Thus, the ability of temporal transcription factor Hunchback to regulate circuit assembly is not limited to one stem cell lineage. Third, we find that the temporal transcription factors, Castor and Pdm, alter synaptic partnerships of NB7-1 neuronal progeny. Thus, multiple temporal transcription factors are able to control circuit membership. Together, these findings provide strong evidence supporting the idea that temporal transcription factors are uniquely potent regulators of decisions that contribute to circuit membership.

**Figure 1.**
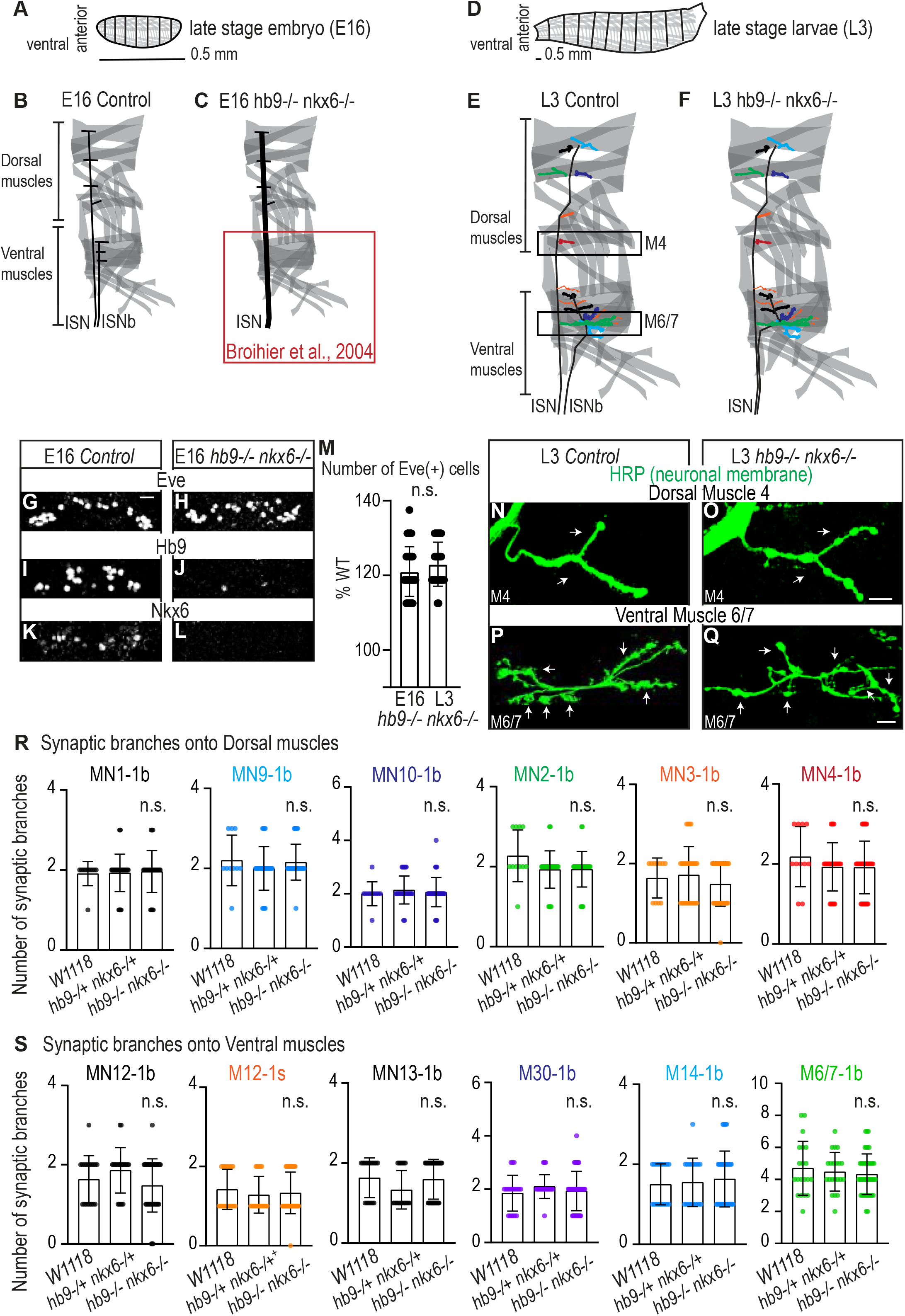
Nk6 Hb9 double mutant does not impact neuromuscular synaptic formation in late stage larvae. (A) Illustration of the late stage embryo, body is organized into repeated left-right, mirror image hemisegments. Muscles (in gray) have a stereotyped pattern. (B-C) Illustration of muscles, ISN nerve that contact dorsal muscles, and ISNb nerve that contact ventral muscles in abdominal bodywall hemisegments at stage 16 in embryos (E16). (C) Illustration representing findings from Broihier et al., 2004. Hb9 nkx6 double mutants lack motor axons of the ISNb nerve and the ISN is thicker than normal. (D) Illustration is the same as in (A), except the body is of a late stage larvae (L3). (E-F) Illustration is the same as in (B-C), except at L3 and synaptic branches are shown. Color code matches axon synapse names in (Q, R). Note, 1s (yellow on ventral muscles) is only scored on Muscle 12 as M12-1s. (E) Illustration of normal synaptic branch pattern at late larval stage (L3). (G) Illustration of L3 synaptic branch pattern seen in hb9 nkx6 mutants, there is no change from normal, wildtype conditions. (G-L) Images of nerve cord segments in stage 16 embryos. In hb9 nkx6 mutants, extra Eve(+) motor neurons are produced while Hb9(+) and Nkx6(+) motor neurons are absent. These findings have been previously published in Broihier et al., 2004. Images are shown anterior up, scale bar represents 15 microns. (M) Quantification of the number of Eve(+) cells represented as a percentage of Control numbers. There is no difference in the number of Eve(+) cells at embryonic stage 16 (E16) or late larval stage (L3). (N-Q) Images of neuronal membrane, both axons and neuromuscular synapses, on ventral muscles in L3 abdominal segments. There is no change in branch number onto each RP muscle target. Data quantified in (I-L). Images are shown dorsal up, anterior left, scale bar represents 10 microns. (R-S) Quantification of the number of 1b or 1s branches on L3 muscles. Color code as in (A) Each dot represents the number of branches onto a single muscle. There is no significant change in synaptic branch number compared to W1118. (R) For w1118 n=11; nkx6−/+, hb9−/+ n=14; nkx6−/−, hb9−/− n=15. (S) For w1118 n=35; nkx6−/+, hb9−/+ n=9; nkx6−/−, hb9−/− n=12. Control is *W1118*. hb9−/− nkx6−/− is *hb9(KK30), Nkx6 (D25)/ hb9(KK30), Nkx6 (D25)*. hb9−/+ nkx6−/+ is *hb9(KK30), Nkx6 (D25)/+*. For quantifications average and standard deviation are overlaid. ANOVA, corrected for multiple samples ‘ns’ not significant.

## Results

### Increasing the number of Eve(+) motor neurons by removal of Nkx6 and Hb9 has no impact on motor neuron-to-muscle synaptic connectivity in larval stages

Prolonged expression of the temporal transcription factor, Hunchback increases the number of Eve(+) motor neurons that make functional synapses onto dorsal muscles at late larval stages (Meng et al., 2019). Eve is a cardinal transcription factor proposed to specify dorsal motor neuron fate including motor axon targeting to dorsal muscles (Labrador et al., 2005). Together this raises the possibility that the cardinal transcription factor Eve is a major downstream effector of Hunchback. If this is the case, then any manipulation that increases the number of Eve(+) motor neurons should lead to an increased number of synaptic branches on dorsal muscles at larval stages. We generate ectopic Eve(+) motor neurons by removing the cardinal transcription factors Nkx6 and Hb9. Nkx6 and Hb9 are normally expressed in motor neurons that project to ventral muscles (Broihier et al., 2002; Broihier et al., 2004). In Nkx6 Hb9 double mutant embryos, Eve repression in ventral motor neurons is lost, leading to an increase in the total number of Eve(+) motor neurons (Broihier et al., 2004). Additionally, in Nkx6 and Hb9 double mutants there is a loss of axons in the ISNb nerve roots, which project to ventral muscles, and increased thickness of the ISN nerve root, which projects to dorsal muscles (Broihier et al., 2004, Figure 1C). Thus, Nkx6 Hb9 double mutants are an excellent genetic background in which to test the hypothesis that increasing Eve(+) motor neurons leads to alterations in muscle innervation in larval stages.

In embryos, we confirmed the Nkx6 and Hb9 double mutant phenotype. We find axon guidance phenotypes, loss of Nkx6 and Hb9 expression (Figure 1I-L), and an increase in Eve(+) neurons (Figure 1G-H). Next, we raised Nkx6 and Hb9 double mutants until late larval stages. The increase in Eve(+) motor neurons in embryos is maintained in late larvae (Figure 1M). We assayed muscle innervation in dissected fillet preparations by visualizing muscles (phalloidin) and neuronal membrane (anti-HRP) and scoring the number of synaptic branches per muscle (Figure 1N-Q). In Nkx6 Hb9 double mutants, the number of synaptic branches onto dorsal or ventral muscles is unchanged compared to wild-type or heterozygous controls (Figure 1F,R-S). Thus, Nkx6 and Hb9 are not required for normal, stable motor neuron-to-ventral muscle partner choice, and Eve expression in motor neurons is not sufficient to generate motor neuron synapses onto dorsal muscles.

In conclusion our data show that changes in the expression of motor neuron cardinal transcription factors Eve, Hb9, and Nkx6 do not lead to permanent alterations in motor neuron-to-muscle synaptic partnerships. This data supports a model where there is an expected pattern of wiring in the Drosophila larval motor system. And, in the case where cardinal transcription factors are manipulated to alter neuronal developmental trajectories, a compensatory program returns the motor system to the expected pattern of wiring. Furthermore, these data show that increasing in the number of Eve(+) motor neurons is not sufficient to explain how prolonged expression of Hb alters circuit membership of NB7-1 progeny. They raise the possibility that temporal transcription factors have a uniquely potent ability to evade compensation, to direct motor neuron-to-muscle synaptic partnerships, and to determine circuit membership of neurons from a given neuroblast.

### Prolonged expression of Hunchback in NB3-1 anatomically alters RP motor neuromuscular synapses

In the Drosophila motor system, temporal transcription factors are expressed in nearly all neuroblasts, where they are thought to control the temporal identity (e.g., first-born, second-born) of neurons. To show that temporal transcription factors, as a class of molecules, control circuit assembly, we need to show that Hunchback can control the circuit membership of neuronal progeny from more than one neuroblast lineage. In this section, we use NB3-1 as a model lineage, and we assay circuit membership using anatomical criteria.

We focus on NB3-1 because temporal transcription factors have been characterized in this lineage, and because NB3-1 produces a group of well-characterized motor neurons (Tran and Doe, 2008). Specifically, NB3-1 generates four Nkx6(+), Hb9(+) ventral motor neurons called RPs in the following birth order: RP1, RP4, RP3, and RP5 (Figure 2A). At larval stages, RP1, RP4, and RP3 project to ventral muscles and make stable Type 1 big (1b) neuromuscular synapses onto Muscles 14, 30 and 6/7, respectively (Landgraf et al., 1997; Choi et al., 2003; Zarin et al., 2019) (Figure 2A, Figure 2--figure supplement 1B). Previous studies, that relied on retrograde Dil labeling, do not agree on the specific ventral muscle target(s) of RP5 (Hoang and Chiba, 2001, Landgraf et al., 1997, Sink and Whitington 1991, Mauss et al., 2009, Schmid et al., 1999). Here, we find that RP5 is a type 1 small (1s) motor neuron (Figure 2--figure supplement 1D). To do so, we used a *Dip-alpha-GAL4*, which drives GFP expression in type 1s motor neurons (herein Dip-alpha>GFP, Ashley et al., 2019; Pérez-Moreno and O’Kane, 2019), and we immunolabel RP1/4/3 (Hb9+, Kr+, Cut-) and RP5 (Hb9+, Kr-, Cut+) (Figure 2--figure supplement 1A, C). We find Dip-alpha>GFP expression specifically in RP5. We conclude RP5 is the ventrally projecting 1s motor neuron and that by late larval stages RP motor neurons from NB3-1 populate the ventral muscle circuit. These findings make NB3-1 a model lineage in which we can test the extent to which Hunchback regulates circuit membership of neurons.

**Figure 2.**
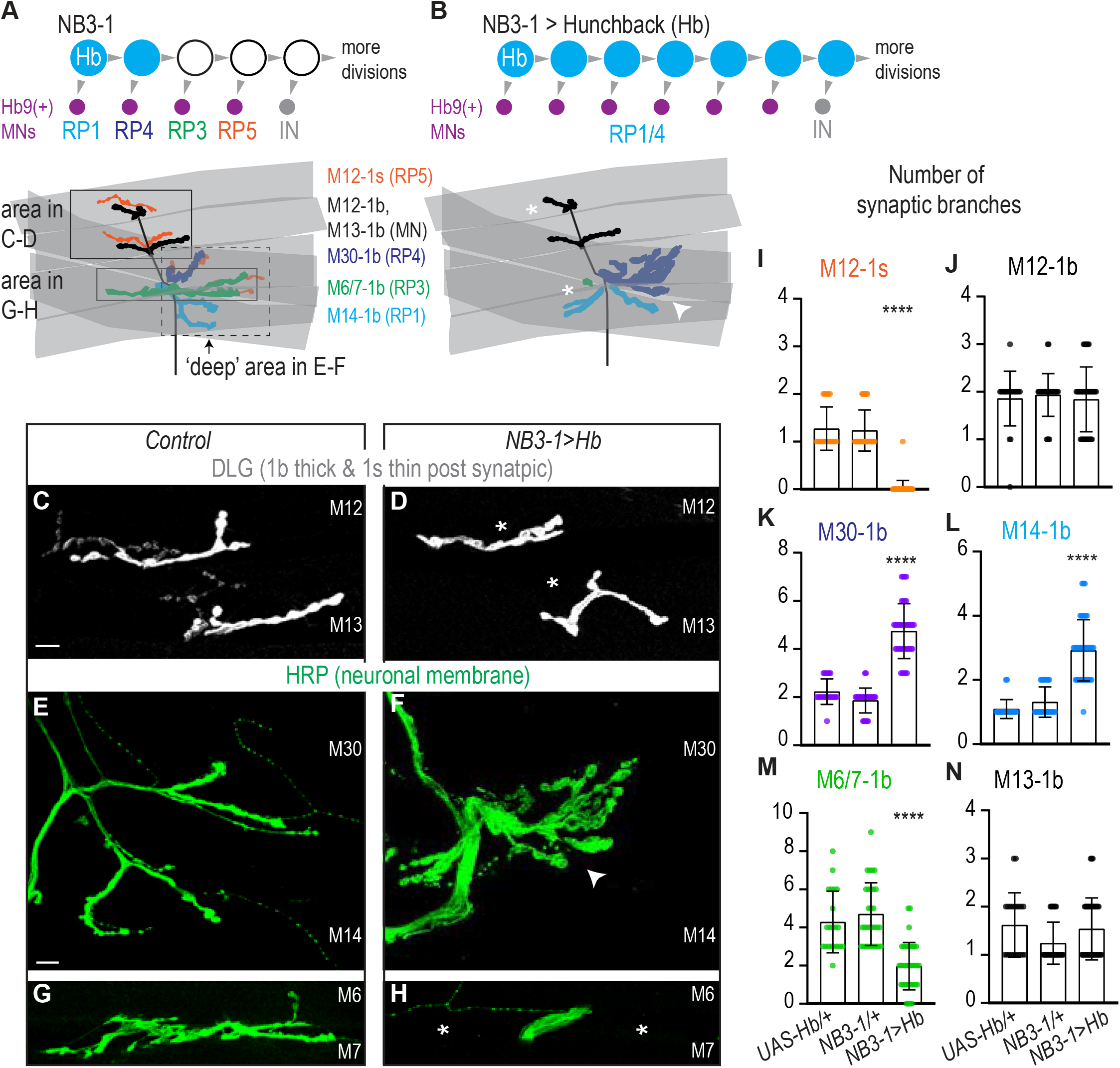
Prolonged expression of Hunchback alters RP motor neuron synapses. (A-B) Illustrations of NB3-1 lineage progression. Each gray arrowhead represents cell division. Each gray arrowhead represents cell division. Large circles are neuroblasts, and smaller circles are neurons. Abbreviations: IN is interneuron. Illustrations of neuromuscular synapses on dorsal muscles in a L3 body wall segment and embryonic molecular identity depicted as circles (blue=RP1/4, green=RP3, orange=RP5). In NB3-1>Hb the number of synaptic branches (in blue) are increased (white arrowhead) onto Muscle 14 and 30 (RP1 and RP4 muscle targets, respectively), 1b synaptic branch number is decreased (asterisk) on Muscle 6 and 7 (RP3 muscle target) and 1s synaptic branch number is lost (asterisk) assessed on Muscle 12 (RP5 muscle target). (C-H) Images of neuronal membrane, both axons and neuromuscular synapses, on ventral muscles in L3 abdominal segments. Arrow indicates branching onto Muscle 30. An asterisk * indicates missing synapses. Data quantified in (I-N). (I-N) Quantification of the number of 1b or 1s branches on L3 muscles. Color code as in (A). Each dot represents the number of branches onto a single muscle. (I-N) For UAS-Hb/+ n=22,21,22,22,21,21. For NB3-1/+ n=30,30,29,29,30,29. For NB3-1>Hb n=39,38,39,39,38,39. Control is *NB3-1/+*. NB3-1>Hb is *NB3-1 GAL4/UAS Hb; UAS Hb/+*. All images are shown dorsal up, anterior left. Scale bars represent 10 microns. For quantifications average and standard deviation are overlaid. ANOVA, corrected for multiple samples ‘****’ p<0.0001.

Next, we prolonged the expression of Hunchback specifically in the NB3-1 lineage (Figure 2--figure supplement 2A-B). To prolong expression of Hb, we drive *UAS-Hunchback* using a NB3-1 specific GAL4 line *“NB3-1-GAL4” (*herein, NB3-1>Hb Figure 2--figure supplement 2C) (Lacin and Truman, 2016). Using a NB3-1 specific driver is critical to rule out lineage non-specific effects, and allows embryos to be healthy enough to grow to larval stages. We confirmed that NB3-1>Hb transforms RP motor neuron identities in a manner consistent with previous work, where Hunchback was overexpressed in all neuroblasts (Tran and Doe, 2008). In NB3-1>Hb embryos, there is an average of six ventral motor neurons (Hb9+, pMad+), that express markers characteristic of early-born RP motor neuron identity (RP1/RP4 Hb+, Kr+, Zfh2-, Cut-), but not later-born RP motor neuron identity (RP3 Hb- Kr+ Zfh2+ Cut- and RP5: Hb-Kr-Zfh2+ Cut+) (Figure 2--figure supplement 2B,D, E). These data confirm that our reagents are working as expected. They show that prolonged expression of Hunchback in NB3-1 increases the total number of ventral motor neurons at embryonic stages.

To determine if, in NB3-1>Hb, increased RP motor neurons at embryonic stages results in permanent anatomical changes to motor circuits, we examined motor neuron-to-ventral muscle synapses in late stage larvae. First, we counted the number of Type 1b and Type 1s synaptic branches on ventral muscles. In wild-type, one type 1b motor neuron makes one or two synaptic branches per muscle (Figure 2A). In NB3-1>Hb, which has increased numbers of RP1/4 motor neurons compared to control, we find a significant increase in the number of type 1b synaptic branch onto Muscle 14 and 30 (RP1 and RP4 muscle targets, respectively)(Figure 2B,E-F,K-L). Additionally, in NB3-1>Hb, which has a decrease in numbers of RP3 and RP5 motor neurons, we see a significant decrease in 1b branch number onto Muscles6/7 (RP3 muscle target) and a significant decrease in 1s branch number onto Muscle 12 (a RP5 muscle target) (Figure 2A-B,C-D,G-H,I-J,M-N). Taken together, we conclude that the increase in the number of embryonic neurons expressing early born RP markers explains the motor neuron-to-muscle partnership choices in the ventral muscle field of late stage larvae.

Next, we examine the extent to which synaptic branches are likely to contain functional synapses (Figure 3B). To do so, we stained for pre-synaptic active zone marker (Bruchpilot), a post-synaptic marker (Discs large), and a post-synaptic neurotransmitter receptor (GluRIIA) (Figure 3A). In NB3-1>Hb, in comparison to controls, we find normal abundance and localization for all markers (Figure 3C-H). These data strongly suggest that the extra synaptic branches on ventral muscles in NB3-1>Hb contain functional synapses.

**Figure 3.**
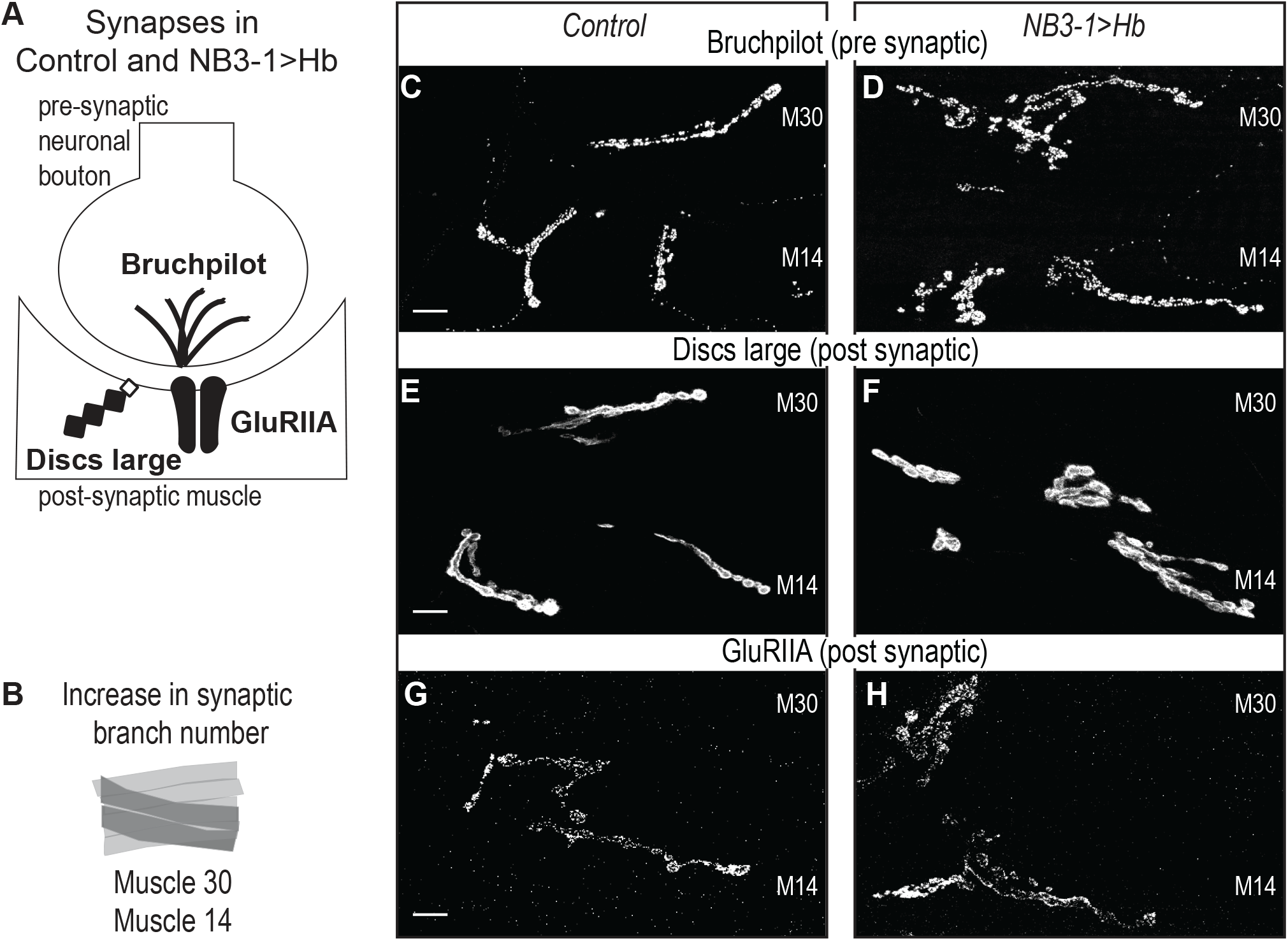
Increased synaptic branches contain pre and postsynaptic markers necessary for function. (A) Illustration of subcellular localization of neuromuscular synapse markers. Brunchpilot labels active zones, Discs large is a scaffolding protein strongly localized at post-synapse, and GluRIIA is the post-synaptic glutamate receptor IIA. (B) Illustration highlighting (darkened) muscles 14 and 30, which have increased synaptic branching in NB3-1>Hb (see Figure 2). (C-H) Images of neuromuscular synapses on L3 Muscle 14 and 30 (see B). There is no difference in distribution or abundance of synaptic markers between Control and NB3-1>Hb. Control is *Hb/+* and NB3-1>Hb is *NB3-1-GAL4/UAS-Hb*; *UAS-Hb/+*. All images are shown dorsal up, anterior to the left, scale bars represent 10 microns.

In summary, we conclude that in wild type animals, NB3-1 produces four RP motor neurons that populate the ventral muscle circuit. Prolonged expression of Hunchback in NB3-1 generates additional RP motor neurons at embryonic stages, all of which have markers for the earliest-born RP motor neuron identities. In NB3-1>Hb, the changes in motor neuron number and embryonic identity are long-lasting because these motor neurons form anatomical synapses on ventral muscle at late larval stages. Our data also support the conclusion that there is an increase in total synapse number on Muscles 14 and 30 and a decrease in total synapse number on Muscles 6 and 7. These data demonstrate that Hunchback can control the anatomical circuit membership of neuronal progeny from more than one neuroblast lineage.

### Prolonged expression of Hunchback in NB3-1 functionally alters RP motor neuromuscular synapses

In this section, we assess to what extent prolonged expression of Hunchback in NB3-1 impacts larval physiology. This is an open question because changing the number of synaptic contacts onto a muscle does not necessarily mean that there will be a corresponding functional change in the muscle. In the Drosophila motor system, there exist well-characterized, homeostatic mechanisms that keep the muscle’s response to nerve stimulation within a set range of values (Frank et al., 2020). This means that anatomical changes do not necessarily lead to functional alterations.

Here, we use electrophysiology to determine the extent to which prolonged expression of Hb in NB3-1 affects motor system physiology. Specifically, we place a stimulating electrode on the nerve root to stimulate release of synaptic vesicles from pre-synaptic motor neurons, and we use a sharp electrode in the muscle to record the post-synaptic response. This provides three measurements: (1) excitatory post-synaptic potential (EPSP), which quantifies the muscles response to nerve stimulation; (2) miniature-EPSP (mEPSP), which quantifies the muscle response to spontaneous release of single synaptic vesicles and is a proxy for postsynaptic receptor sensitivity; and (3) quantal content (EPSP/mEPSP), which estimates the numbers of vesicles released during nerve stimulation (Figure 4A).

**Figure 4.**
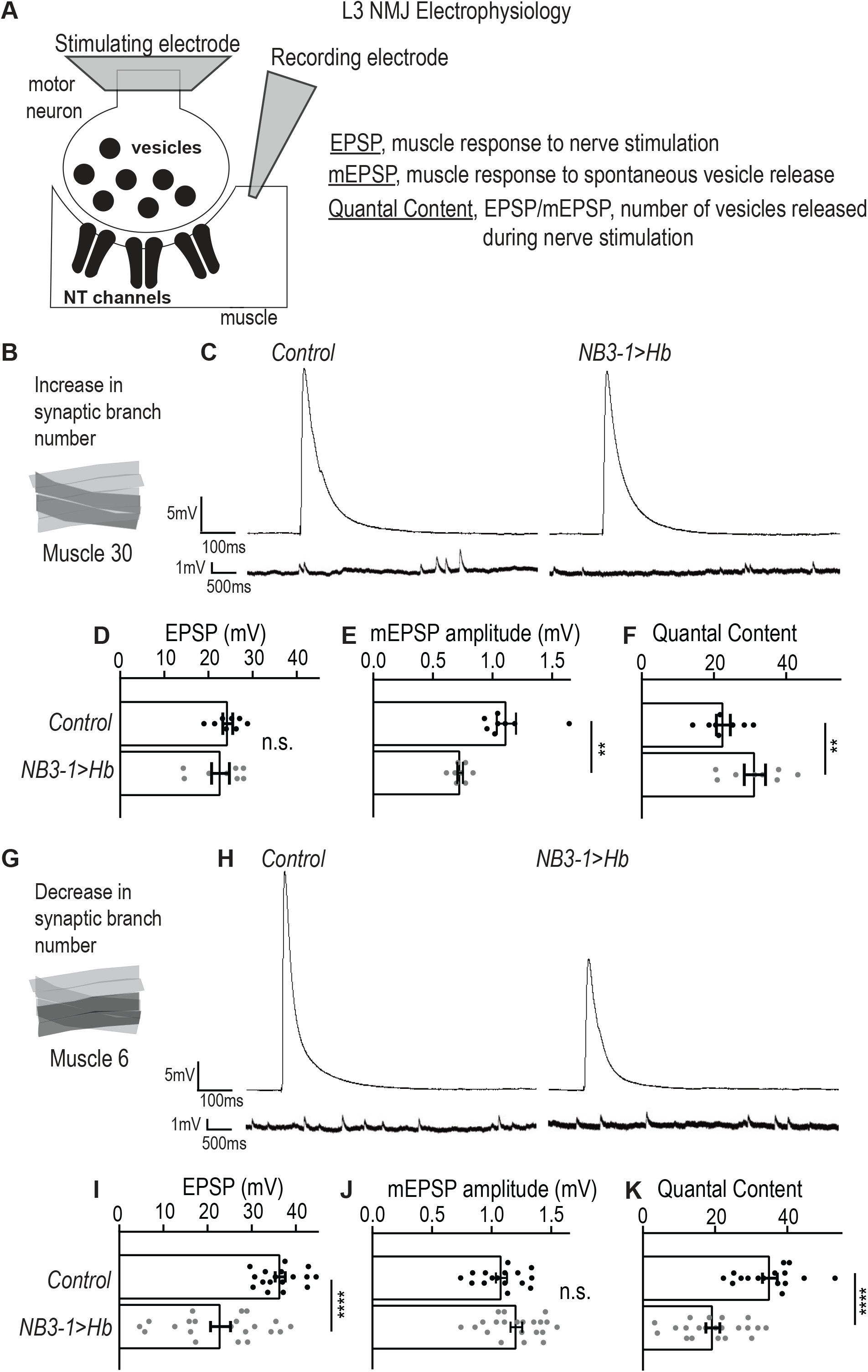
Altered synapses onto Ventral muscles are functional. (A) Illustration of neuromuscular junction (NMJ) electrophysiology approach on third instar larvae (L3). Abbreviation NT is neurotransmitter. (B) Illustration highlighting (darkened) Muscle 30 whose electrophysiological recordings are presented in C-F. (C) Traces of EPSP and mEPSPs for Muscle 30. (D-F) Quantification of electrophysiological recordings for Muscle 30. (D) Evoked response (EPSP) is not changed in Muscle 30 in NB3-1>Hb vs Control. (E) Spontaneous response (mEPSP) is significantly decreased in Muscle 30 in NB3-1 vs Control. (F) Transmitter release (Quantal Content (EPSP/mEPSP)) is slightly significantly increased in Muscle 30 NB3-1 vs Control. (D-F) For Control, n= 8. For NB3-1>Hb, n=8. For C-E and H-J, each dot (black represents Control and gray represents NB3-1>Hb) on the graph represents a single recording from a unique bodywall segment from A2-A4. (G) Illustration highlighting (darkened) Muscle 6 whose electrophysiological recordings are presented in H-K. (H) Traces of EPSP and mEPSPs for Muscle 6. (I-K) Quantification of electrophysiological recordings for Muscle 6. (I) Evoked response (EPSP) is significantly decreased in Muscle 6 in NB3-1>Hb vs Control. (J) Spontaneous response (mEPSP) is not changed in Muscle 6 in NB3-1 vs Control. (K) Transmitter release (Quantal Content (EPSP/mEPSP)) is significantly decreased in Muscle 6 NB3-1 vs Control. (I-K) For Control n= 17,16,16. For NB3-1>Hb n=21. Control is NB3-1/+ and NB3-1>Hb is NB3-1-GAL4/UAS-Hb; UAS-Hb/+. All images are shown dorsal up, anterior to the left. For quantifications average and standard deviation are overlaid. Unpaired t-test ‘ns’ not significant, ‘**’ p<0.05, ‘****’ p<0.0001.

First, we examined the physiology of Muscle 30. In NB3-1>Hb, there is an increase in nerve branches and synaptic markers on Muscle 30 (Figure 4B). Increases in synapse number on a muscle is expected to increase the response of the muscle to nerve stimulation. However, we observe no increase in EPSP amplitude in NB3-1>Hb, compared to wild type (Figure 4C-D). This suggests homeostasis alters the system to maintain the muscle response to nerve stimulation. Conceptually there are two ways homeostasis could be working in this system: 1) neurotransmitter receptors on the postsynaptic muscle could be less sensitive to neurotransmitters (Frank et al., 2006); 2) synapses might release fewer vesicles upon nerve stimulation. To examine these possibilities, we measured mEPSP and quantal content. In NB3-1>Hb, there is a significant increase in quantal content when compared to control animals (Figure 4C,E-F). This suggests that upon nerve root stimulation, the number of presynaptic vesicles released is increased, consistent with the idea that the total number of synapses is increased in NB3-1>Hb. However, in NB3-1>Hb, there is a significant decrease in mEPSP amplitude compared to control animals (Figure 4C,E). These data suggest that the postsynaptic receptor sensitivity to neurotransmitter is reduced.

Next, we examined the physiology of Muscle 6, which has a decrease in synaptic branches in NB3-1>Hb (Figure 4G). If there is no homeostatic compensation, then decreased number of synapses is expected to decrease the presynaptic neurotransmitter release (quantal content), not affect postsynaptic receptor sensitivity (mEPSP amplitude), and lead to an overall decrease in the muscle response to nerve stimulation (EPSP amplitude) (Figure 4H,I-K). Our data are consistent with these expectations. Together, our results suggest an impaired synaptic function and lack of homeostatic compensation on Muscle 6. Similar phenotypes have been observed in previous studies, where reduced synaptic number is correlated with decreased synaptic function (Goel et al., 2019).

We conclude that in NB3-1>Hb, on Muscle 30, wild type levels of synaptic function are maintained. Whereas, in NB3-1>Hb, Muscle 6 synaptic function is impaired. Taken together these data show that prolonged expression of Hunchback in NB3-1 results in functional changes in motor system physiology.

### Precocious expression of Castor in NB7-1 decreases the number of U motor neurons and causes loss of synaptic connectivity onto later born U muscle targets

In the Drosophila motor system, Hunchback is one of several temporal transcription factors in a well-characterized cascade, which also includes Kruppel, Pdm, and Cas. To show that temporal transcription factors as a class of molecules regulate circuit membership, we need to manipulate temporal transcription factors other than Hunchback. For the remainder of this paper, we focus on two temporal transcription factors, Castor and Pdm. We focus on Castor and Pdm because, although they are temporal transcription factors like Hunchback, they differ from Hunchback in important ways. First, Hunchback is the earliest transcription factor in the series, whereas Castor and Pdm are expressed later in the series. Second, Hunchback is thought to specify early-born temporal identities, whereas Castor and Pdm specify later-born temporal identities. Third, prolonged expression of Hunchback extends the time that neuroblasts produce early-born neurons, whereas precocious expression of Castor and Pdm can truncate the time that neuroblasts produce early-born neurons. Together the series of experiments described in the sections below probe the extent to which changes in temporal transcription factors Cas and Pdm produce long lasting changes in synaptic connectivity and thus, circuit membership.

Here, we use NB7-1 as a model lineage because temporal transcription factors are arguably the most well-characterized in the NB7-1 lineage. Early divisions of NB7-1 give rise to five U motor neurons in the following birth order: U1, U2, U3, U4, and U5 (Figure 5A). U motor neurons all express the cardinal transcription factor Eve and all U motor neurons populate the dorsal muscle circuit (Figure 5A). In embryos, each U motor neuron can be identified by a unique combination of marker genes, and in larvae, each U motor neuron synapses on a unique dorsal muscle (Figure 5--figure supplement 1A). Studying NB7-1 as a model lineage allows us to investigate the impact of manipulating temporal transcription factors on embryonic temporal identities and to directly link this to mature synaptic partnerships.

**Figure 5.**
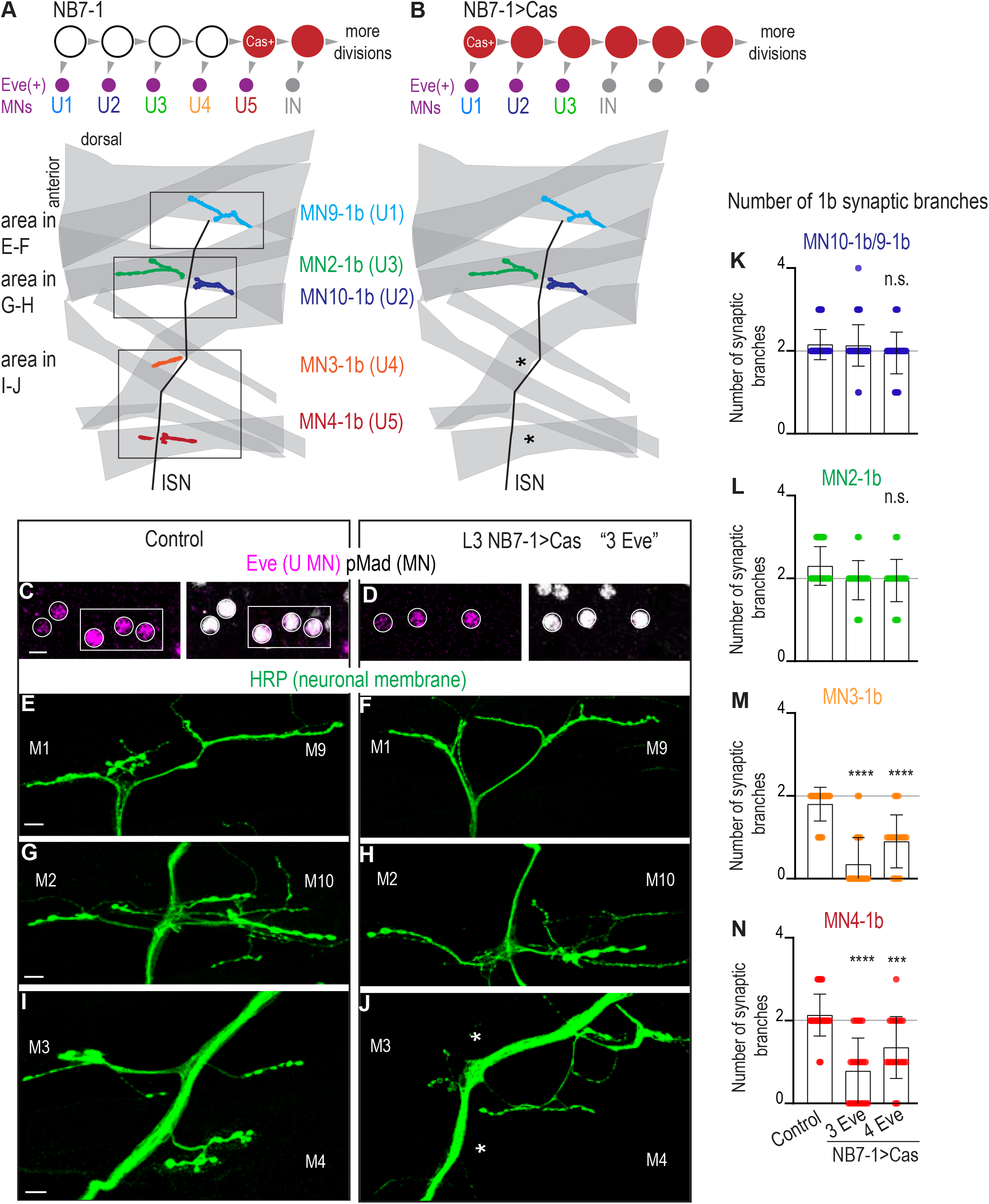
Precocious expression of Castor alters U motor neuron synapses. (A-B) Illustrations of NB7-1 lineage progression. Each gray arrowhead represents cell division. Each gray arrowhead represents cell division. Large circles are neuroblasts, and smaller circles are neurons. IN is interneuron. In NB7-1>Cas there is a decrease in the number of Eve(+) neurons with U4/U5 embryonic molecular identity. Illustrations of neuromuscular synapses on dorsal muscles in a L3 body wall segment and embryonic molecular identity depicted as circles (light blue=U1, dark blue=U2, green=U3, orange=U4, red=U5). (C-D) Images of L3 nerve cord abdominal hemisegment Eve(+) neurons co-expressing the motor neuron marker pMad. For Control, n=12 hemisegments in 3 animals. For NB7-1>Cas, n=23 hemisegments in 7 animals. All images are shown anterior up, midline left, scale bar represents 5 microns. (E-J) Images of neuronal membrane, both axons and neuromuscular synapses, on ventral muscles in L3 abdominal segments corresponding to hemisegments containing the number of Eve(+) neurons as imaged in (D-E). An asterisk * indicates missing synapses on Muscle 3 and Muscle 4. All images are shown dorsal up, anterior left, scale bar represents 10 microns. Data quantified in (K-N). (K-N) Quantification of the number of 1b branches on L3 muscles. Color code as in (A). Line intersects the y-axis at 2. Each dot represents the number of branches onto a single muscle. Decrease in synaptic branching onto Muscle 4 and Muscle 3 in experimental conditions (3 Eve and 4 Eve) vs Control (K). No change (L). (K-N) For Control n=59,30,30,30. For NB7-1>Cas 3 Eve n=46,23,23,23. For NB7-1>Cas 4 Eve n=40,20,20,20. Control is *Cas/+* and NB7-1>Cas is *NB7-1 GAL4/Cas*. For quantifications, average and standard deviation are overlaid. ANOVA, corrected for multiple samples ‘ns’ not significant, ‘***’ p<0.001, ‘****’ p<0.0001.

Before examining larval synapses, we confirmed that our Cas manipulation generates embryonic motor neuron phenotypes. In wild type, in NB7-1, Cas is first expressed during the division that specifies the temporal identity associated with U5, and Cas continues to be expressed in NB7-1 during subsequent divisions that give rise to a set of Cas(+) interneurons (Figure 5--figure supplement 1A). Cas expression closes the window of U motor neuron production. This can be seen by driving UAS-Cas with En-GAL4, which is expressed from the first division in all nerve cord row six and seven neuroblasts, and observing the production of fewer than five U motor neurons (Grosskortenhaus et al., 2006). Here, we express Castor starting during the earliest divisions of NB7-1 using a NB7-1 specific driver, *NB7-1-GAL4* to drive *UAS-Cas* (herein, NB7-1>Cas)(Figure 5--figure supplement 1B). NB7-1>Cas allows us to grow animals to late larval stages and to rule out any non-lineage specific effects. We stained NB7-1>Cas embryos with a U motor neuron marker (Eve), a pan-motor neuron marker (pMad), and markers for individual U motor neuron identities (Figure 5--figure supplement 1F). In a majority of hemisegments in NB7-1>Cas, we see decreases to sets of three or four U motor neurons (Figure 5--figure supplement 1D). These U motor neuron sets have molecular identities of [U1, U2, U3] or [U1, U2, U3, U4/U5], respectively (Figure 5--figure supplement 1E). These data shows our reagents are working as expected. We conclude that at embryonic stages, precocious expression of Cas, specifically in NB7-1, decreases the number of U motor neurons produced by NB7-1.

Next, in NB7-1>Cas, we characterize mature neuromuscular synapses at larval stages. Because NB7-1>Cas has different numbers of U motor neurons in different hemisegments, we matched nerve cord hemisegments to corresponding body wall segments (Figure 5C-D). In hemisegments in NB7-1>Cas with fewer U motor neurons (three or four), we counted mature synaptic branches onto individual dorsal muscles. Compared to Control, we find no change in 1b branch number onto Muscles 9, 10, and 2 (U1, U2, and U3 muscle targets, respectively) (Figure 5E-H,K-L). However, compared to our control, we see a significant decrease in 1b branch number onto Muscles 3 and 4 (U4 and U5 muscle targets, respectively) (Figure 5I-J,M-N). These data suggest that precocious expression of Cas in NB7-1 leads to changes in embryonic U motor neuron number, and that these changes correspond to long-lasting changes in circuit wiring at larval stages. We conclude that at least one temporal transcription factor other than Hunchback can control the number of motor neurons that populate the dorsal muscle circuit.

### Precocious expression of Pdm in NB7-1 anatomically and functionally alters U motor neuromuscular synapses

Here, we continue to address the question to what extent do temporal transcription factors other than Hb control the number of NB7-1 neuronal progeny that populate the dorsal muscle circuit. Specifically, we manipulate the temporal transcription factor Pdm to generate “heterochronic” mismatches between a U motor neuron’s embryonic temporal identity and its birth time. By studying heterochronic mismatches we can assess the relative contribution of intrinsic factors (e.g., neuronal gene expression) versus extrinsic factors (e.g., availability of synaptic partners or transient signaling cues) in control of synaptic partner selection.

Prolonged Hb expression in NB7-1 generates a heterochronic mismatch. It produces a large number of ectopic motor neurons with U1-like molecular identities (Isshiki et al., 2001). This means that in NB7-1>Hb, <u>*early-born*</u> U motor neurons are generated at <u>*abnormally late*</u> times in development. In NB7-1>Hb, at late larval stages there are increases in motor neurons innervating the dorsal muscles (Meng et al., 2019). This demonstrates that prolonged expression of Hb causes early-born U motor neurons to populate the dorsal muscle without regard to time-linked environmental cues. Notably, precocious expression of the temporal transcription factor Cas in NB7-1, described in the section above, reduces the total number of U motor neurons, but does not create a heterochronic mismatch. Here, using Pdm, we produce two heterochronic mismatches that allow us to determine the extent to which <u>*late-born*</u> U motor neurons born either at <u>*abnormally late*</u> or <u>*abnormally early*</u> times in development can populate dorsal muscle circuits. These experiments are critical for determining the extent to which factors other than Hb provide information to specify synaptic partner selection and circuit membership.

We manipulate Pdm, which is expressed during the fourth and fifth division window of NB7-1. Pdm specifies the U4 temporal identity and represses Kruppel, the temporal transcription factor that specifies the U3 temporal identity (Figure 6A) (Grosskortenhaus et al., 2006). We precociously express Pdm in the NB7-1 using *NB7-1-GAL4* to drive *UAS-Pdm* (herein, NB7-1>Pdm), and characterize this manipulation by immunostaining with markers in embryos for U motor neurons (Eve), all motor neurons (pMad) and individual U motor neuron temporal identity markers (Figure 6--figure supplement 1). We find that NB7-1>Pdm transforms U motor neuron identities in a manner consistent with other GAL4 drivers (Grosskortenhaus et al., 2006). Specifically, in NB7-1>Pdm, we see two phenotypes--either decreases or increases in the number of U motor neurons per abdominal hemisegment in comparison to controls (Figure 6B-C top). Each phenotype is discussed in a separate paragraph below.

**Figure 6.**
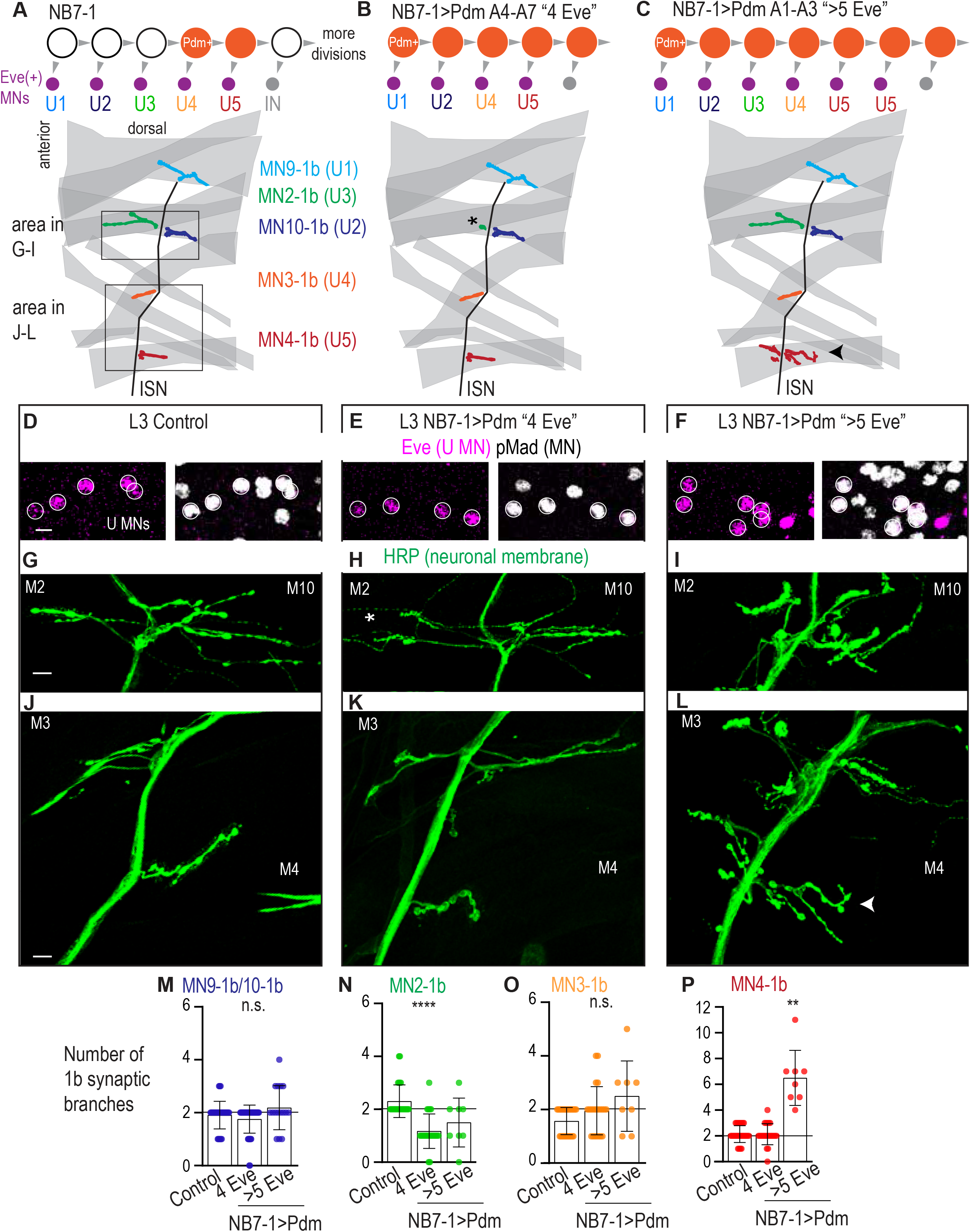
Precocious expression of Pdm alters U motor neuron synapses. (A-C) Illustration of NB7-1 lineage progression. Each gray arrowhead represents cell division. Each gray arrowhead represents cell division. Large circles are neuroblasts, and smaller circles are neurons. Abbreviations: IN is interneuron. Illustrations of neuromuscular synapses on dorsal muscles in a L3 body wall segment and embryonic molecular identity depicted as circles (light blue=U1, dark blue=U2, green=U3, orange=U4, red=U5). (D-F) Images of L3 nerve cord abdominal hemisegment Eve(+) neurons co-expressing the motor neuron marker pMad. All images are shown anterior up, midline left. Scale bars represent 5 microns. (G-L) Images of neuronal membrane, both axons and neuromuscular synapses, on ventral muscles in L3 abdominal segments corresponding to hemisegments containing the number of Eve(+) neurons as imaged in (D-E). An asterisk * indicates missing synapses on Muscle 2 and an arrow indicates increase in synapses on Muscle 4. Data quantified in (M-P). All images are shown dorsal up, anterior left. Scale bars represent 10 microns. (M-P) Quantification of the number of 1b branches on L3 muscles. Color code as in (A). Line intersects the y-axis at 2. Each dot represents the number of branches onto a single muscle. Control is *Pdm/+*. NB7-1>Pdm is *NB7-1 GAL4/Pdm; Pdm/+*. In H, Control is *NB7-1 GAL4/UAS myr GFP*. (M-P) For Control, n=33,26,28,28. For NB7-1>Pdm with 4 Eve(+), n=45,23,22,23. For NB7-1>Pdm with 5 Eve(+), n=16,8,8,8. For quantifications average and standard deviation are overlaid. ANOVA, corrected for multiple samples ‘ns’ not significant, ‘**’ p<0.05, ‘****’ p<0.0001.

In NB7-1>Pdm, hemisegments with additional U motor neurons are examples of heterochronic mismatches in which <u>*later-born*</u> U motor neurons are born at <u>*abnormally late*</u> times in development. Notably, we observe this phenotype selectively in segments A1-A3 (Figure 6C). In these segments, U motor neurons have the following embryonic molecular identities: U1, U2, U3, U4, followed by a variable number of U5-like neurons (n=12/154) (Figure 6C, Figure 6--figure supplement 1E). Therefore, in NB7-1>Pdm, neurons with U5-like identities are born at abnormally late times, i.e., when interneurons are usually made.

In NB7-1>Pdm, hemisegments with fewer U motor neurons are examples of heterochronic mismatches in which <u>*later-born*</u> U motor neurons are born at <u>*abnormally early*</u> times in development. Notably, we observe this phenotype selectively in segments A4-A7 (Figure 6B). In these segments, U motor neurons have the following embryonic molecular identities: U1, U2, U4, and U5, but not U3 (Figure 6C, Figure 6--figure supplement 1E). To confirm this heterochronic mismatch, we co-stained for the U4 and U5 marker Runt and the U motor neuron marker Eve in stage l11/e12 embryos. At this stage in wild type, four Eve(+) neurons are generated, <u>*one*</u> of which is Runt(+). In NB7-1>Pdm, at the same stage, of the four Eve(+) neurons <u>*two*</u> can be Runt(+) (Figure 6--figure supplement 1H). Therefore, in NB7-1>Pdm, NB7-1 can skip production of U3 and produce neurons with U4 and U5 molecular identities at abnormally early times in development.

To determine how NB7-1>Pdm affects mature motor circuits, we looked at motor neuron-to-dorsal muscle synapses in late stage larvae. First, we looked at neuroanatomy, matching nerve cord hemisegments to corresponding body wall segments, and staining for muscle (phalloidin) and neuronal membrane (anti-HRP) (Figure 6A-F). When hemisegments contain >5 Eve(+) pMad(+) neurons (e.g., those with U5-like neurons born at abnormally late times), we find a significant increase in 1b branch number selectively Muscle 4 (i.e., wild type U5 target), while all other muscles had no change in branch number compared to our control (Figure 6A,C,G,I,J,L,M-P). When hemisegments contain 4 Eve(+) pMad(+) neurons (e.g., those with U4 and U5 generated at abnormally early time and lacking a U3-like neuron), compared to our control, we find a significant decrease in 1b branch number on Muscle 2 (U3 muscle target), and no changes in 1b branch number on other muscles (Figure 6A-B,G-H, J-K,M-P). These data show that the alterations in embryonic motor neuron markers in NB7-1>Pdm correlates with changes in the number of synaptic branches on dorsal muscles at larval stages.

Next, we confirmed that in NB7-1>Pdm in abdominal segments A1-A3, the increased nerve branches on Muscle 4 contained neuromuscular synapses. We stained for synaptic proteins for active zones (Bruchpilot), postsynaptic densities (Discs large), and neurotransmitter receptors (GluRIIA) (Figure 7A). We find a normal abundance and localization for all markers (Figure 7B-G). Thus, on the cell biological level, neuromuscular synapses in NB7-1>Pdm are not different from Control.

**Figure 7.**
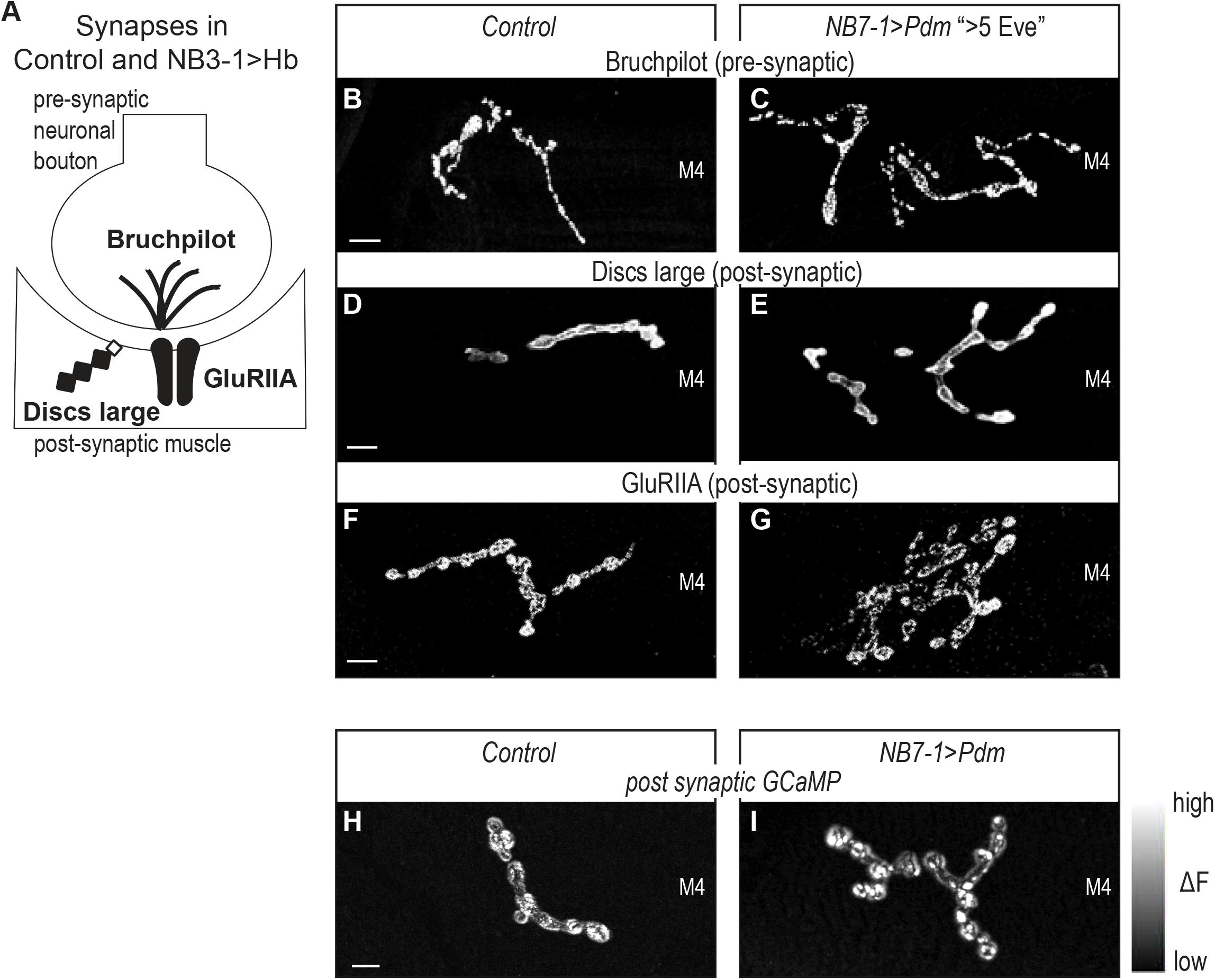
Altered synapses onto Dorsal muscles are functional. (A) Illustration of subcellular localization of neuromuscular synapse markers. Brunchpilot labels active zones, Discs large is a scaffolding protein strongly localized at post-synapse, GluRIIA is a glutamate receptor IIA. (B-G) Images of neuromuscular synapses on L3 Muscle 4. There is no difference in distribution or abundance of synaptic markers between Control and NB7-1>Pdm. Control is *Pdm/+* and NB7-1>Pdm is *NB7-1-GAL4/UAS-Pdm*; *UAS-Pdm/+* (H-I) Images of fluorescence intensity changes in a calcium indicator of synaptic activity. GCaMP was targeted to the post-synaptic density for example (DLG in E-G). When pre-synaptic vesicles are released from active zones (Brp in B-C) post-synaptic neurotransmitter receptors respond (GluRIIA in G-F), increasing GCaMP fluorescence intensity (see Figure 9—figure supplement 1 for details). Images show post-synaptic responses (delta F) in L3 Muscle 4 (M4) in Control and NB7-1>Pdm. Control is *Pdm/+; MHC-CD8-GCaMP6f-Sh/Pdm* and NB7-1>Pdm is *NB7-1-GAL4/Pdm; MHC-CD8-GCaMP6f-Sh/Pdm*. All images are shown dorsal up, anterior to the left. Scale bars represent 10 microns.

Finally, for NB7-1>Pdm (A1-A3), we visualize neuromuscular synapse activity onto Muscle 4 with a post-synaptically localized calcium sensor (Newman et al., 2017). In late larval neuromuscular synapses, we imaged spontaneous release of individual synaptic vesicles from 1b branches on Muscle 4, that have increased numbers of synaptic branches. In both Control and NB7-1>Pdm, we see that within a five-minute imaging period, there is at least one postsynaptic response (Figure 7H-I, Figure 7--figure supplement 1). From these findings, we conclude that Pdm can impact functional synaptic connections onto muscles made by U motor neurons.

In summary, we find that precocious expression of Pdm in NB7-1 leads to two different types of heterochronic mismatches: one in which <u>*later-born*</u> U motor neurons are born at <u>*abnormally late*</u> times in development, and one in which later-born U motor neurons are born at <u>*abnormally early*</u> times in development. We find a strong correlation between embryonic molecular identity and larval neuromuscular synapses. Together these data support the idea that regardless of birth time, both early-born and later-born U motor neurons can populate dorsal muscle circuits. They demonstrate that Hb is not the only temporal transcription that can provide enough molecular information to neuronal progeny, such the neuronal progeny are insensitive to time-linked environmental cues. Ultimately this supports the idea that a general feature of the temporal transcription factors is to control circuit membership.

## Discussion

In this study, we address the question to what extent is control of circuit membership a general feature of temporal transcription factors. Our study has three key findings. First, we show that although deletion of cardinal transcription factors Nkx6 and Hb9 alters motor neuron gene expression and axon guidance, this does not lead to permanent alterations in motor neuron-to-muscle connectivity (Figure 1, Figure 8B). Thus, not all manipulations that alter axon guidance lead to alterations in mature synapses and circuit membership. Next, we find prolonged expression of the temporal transcription factor, Hunchback increases the number of RP motor neurons produced by NB3-1, and this causes permanent changes in the anatomy and function of the ventral muscle circuit (Figure 2-4, 8C). Because we previously showed that Hunchback acts similarly in another neuroblast, NB7-1, we conclude that a general feature of this temporal transcription factor is to control circuit membership of neurons in a lineage (Meng et al., 2019). Third, we show precocious expression of additional temporal transcription factors, Castor and Pdm, alter the number of U motor neurons produced by NB7-1, and that this results in permanent changes in the dorsal muscle circuit (Figure 5-7, 8C-D). Together these findings provide strong support for the hypothesis that temporal transcription factors, as a class of molecules, are key determinants of motor circuit membership.

**Figure 8.**
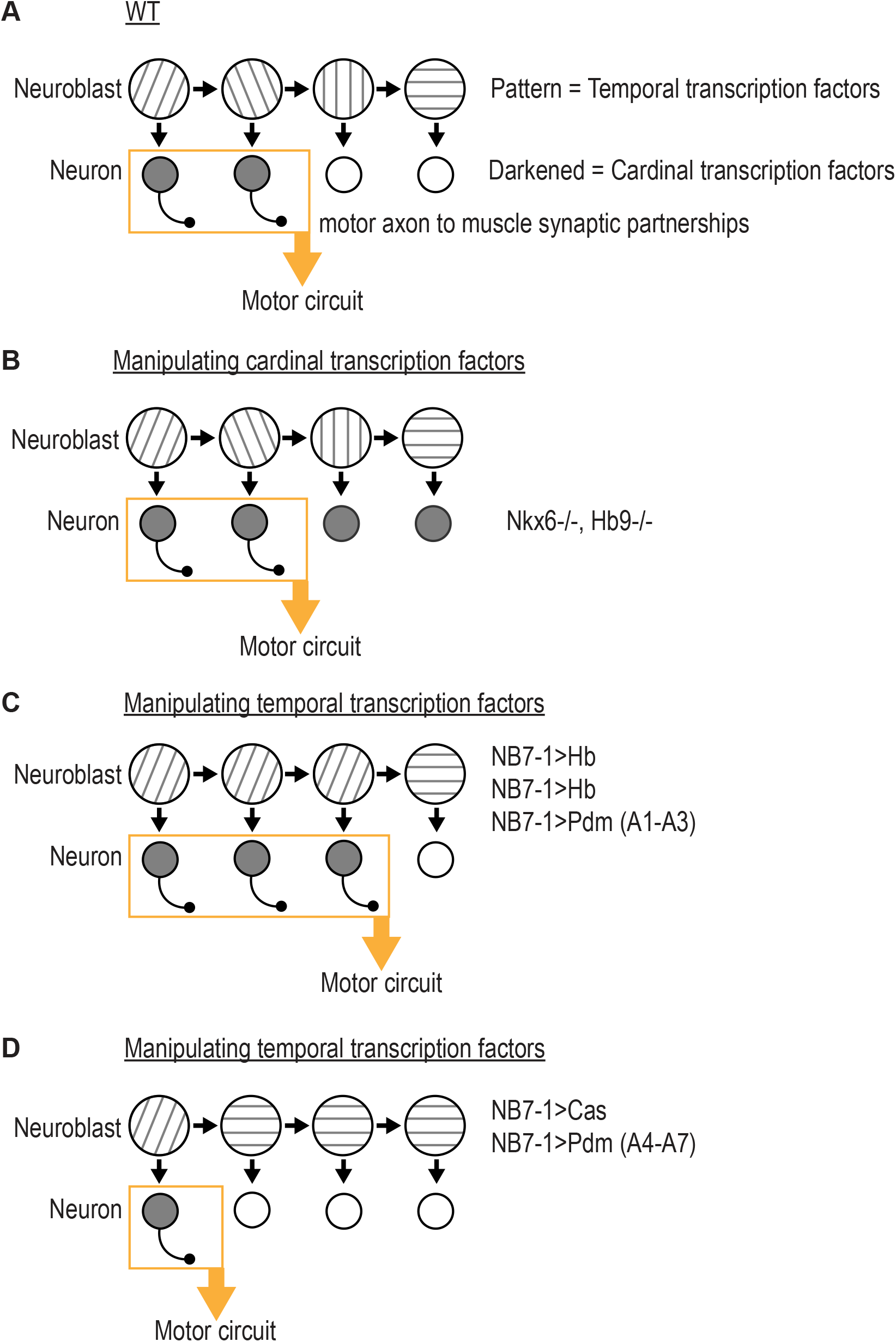
Summary of results in this study. (A) Illustration of WT(wildtype) neuroblast lineage progression. Different patterns in the neuroblast represent temporal transcription factor expression. Darkened neurons represent cardinal transcription factor expression. Outgrowth projecting from neuron represent motor axon to muscle synaptic partnerships. Yellow box and arrow represent that these neurons are members of the same motor circuit. (B) Illustration of the outcome when expression of Nkx6 and Hb9 are lost. Motor axon to muscle synaptic partnerships remain unaltered. (C) Illustration of the outcome when expression of Hb (Hunchback) is prolonged in NB7-1 or NB3-1, or expression of Pdm is precocious in NB7-1 A1-A3 (abdominal segments 1 through 3). Motor axon to muscle synaptic partnerships are altered by increased circuit membership. (D) Illustration of the outcome when expression of Cas (Castor) is precocious in NB7-1 or expression of Pdm is precocious in NB7-1 A4-A7 (abdominal segments 4 through 7). Motor axon to muscle synaptic partnerships are altered by decreased circuit membership.

### Alterations in axonal trajectory can be compensated for later in development

In the Drosophila motor system, cardinal transcription factors such as Eve, Nkx6, and Hb9 are necessary and sufficient to control motor neuron axon outgrowth to particular muscle fields (Seroka et al., 2020 biorxiv; Landgraf et al., 1999; Broihier et al., 2002; Broihier et al., 2004). However, the extent to which early developmental defects lead to long-lasting defects in the formation of motor neuron-to-muscle synaptic partnership was unknown (Landgraf et al., 1999; Fuijoka et al., 2002; Broihier et al., 2004; Broihier et al., 2002). We find in nkx6 and hb9 double mutants, neuromuscular innervation patterns are indistinguishable from wild type by late larval stages (Figure 1). This demonstrates that changes in axonal trajectory in embryos are not sufficient to permanently alter motor neuron axon to muscle synaptic partnerships in Drosophila locomotor circuits (Figure 8A-B). Because embryonic developmental defects are corrected by late larval stages, this suggests that the motor system compensates for the early defect.

The idea that the Drosophila motor system can compensate for early developmental defects in nkx6 hb9 double mutants is consistent with other studies. For example, manipulation of neuronal cell surface molecules such as Connectin and Receptor Tyrosine Phosphatases generate early motor axon guidance defects that ultimately make normal motor neuron-to-muscle synapses, but with a delay (Nose et al., 1994; Desai et al., 1996). In another example, ectopic, pan-neuronal expression of the cardinal transcription factor Eve generates Eve(+) RP3 motor neurons. The Eve(+) RP3 motor neuron axons are routed from their normal ventral muscle targets to the dorsal muscle field (Landgraf et al., 1999). Yet, in some young, pre-hatching larvae, Eve(+) RP3 motor neurons continue to grow from the dorsal muscle region back to the ventral muscles, where they form synapses with their normal RP3 synaptic partners (ventral muscles 6/7) (Landgraf et al., 1999). We speculate that this phenotype may be an intermediate, and if it was possible to rear these animals to late larval stages, they would have wild type muscle innervation patterns. Together with our data, these studies suggest that multiple different early perturbations in the Drosophila motor system can be corrected by late larval stages.

The idea that compensation exists in the Drosophila motor system suggests the following model: First, there is an expected pattern of motor neuron-to-muscle synaptic partnerships. Second, deviations from this expected pattern can be detected. Third, deviations can be corrected. Such a model raises a number of questions: What types of perturbations can be corrected? How are perturbations detected? What is the nature of the compensatory program(s) that correct early defects?

In this study, we show that manipulation of temporal transcription factors permanently alters motor neuron-to-muscle connectivity in Drosophila locomotor circuits. This observation suggests that perturbations induced by temporal transcription factors do not get corrected, and so compensation is limited. We speculate that this limitation may be tied to the timing and scale of the perturbation during development. For example, temporal transcription factors act in neuroblasts whereas cardinal transcription factors act in post-mitotic neurons. Additionally, temporal transcription factors are likely to regulate a large gene network, and a subset of genes within this network are cardinal transcription factors. Cardinal transcription factors themselves are likely to regulate a smaller gene network. Therefore, temporal transcription factors may evade compensation because they are involved in setting up the expected pattern of motor neuron-to-muscle synaptic partnerships. Future work is needed to test such a model.

### A general feature of the temporal transcription factor Hunchback is to control circuit membership of neurons in a lineage

In Drosophila motor system development, most, if not all, neuroblasts express the temporal transcription factor Hunchback early during lineage development (Pearson and Doe, 2004). In all lineages tested so far, Hunchback is necessary and sufficient to specify early-born identity (e.g., first-born), regardless of the cell type (e.g., motor neuron, interneuron, glia) (Cleary and Doe, 2006; Isshiki et al., 2001; Kambadur et al., 1998; Novotny et al., 2002; Pearson and Doe, 2003). Thus, Hunchback is considered to be a temporal transcription factor, which is a class of transcription factor postulated to control entire time-linked developmental programs. Notably, the mammalian homolog of Hunchback, Ikaros, acts as a temporal transcription factor in both retina and cortex (Alsiö et al., 2013; Elliott et al., 2008). In support of the idea that Hunchback controls time-linked developmental programs, Hunchback has been shown to regulate many aspects of time linked post-mitotic neuronal identity, including cardinal transcription factor expression (e.g., Hb9 and Eve) and embryonic axonal trajectories (Isshiki et al., 2001; Tran and Doe, 2008; Seroka and Doe, 2019; Meng et al., 2019). However, until recently, how Hunchback-induced changes in early neuronal development impacted later stages of circuit assembly was unknown.

A pair of recent studies from Seroka and Meng sought to understand the extent to which Hunchback played a role in controlling later aspects of neuronal development (Seroka et al., 2019; Meng et al., 2019). They showed that prolonging the expression of Hunchback in NB7-1 dramatically increases the number of Eve(+) ectopic U motor neurons produced by this neuroblast, and that ectopic U motor neurons project axons to dorsal muscles (Seroka et al., 2019; Meng et al., 2019). Meng et al went on to show that ectopic U motor neurons form permanent functional synapses on dorsal muscles in the dorsal motor circuit (Meng et al., 2019). Therefore, in NB7-1, Hunchback acts as a molecular switch that determines what proportion of neurons from the lineage populate the dorsal muscle circuit.

Here, we addressed the question, to what extent is control of circuit membership a general feature of Hunchback. We prolonged Hunchback in NB3-1 neuroblast, which produces a series of ventrally projecting, Hb9(+) RP motor neurons (Tran and Doe, 2008). Notably, NB3-1 begins dividing much later in nerve cord development than NB7-1 (Bossing et al., 1996). Prolonging the expression of Hunchback in NB3-1 dramatically increased the number of Hb9(+) motor neurons produced and caused permanent changes in neuromuscular synaptic partnerships and physiology in the ventral muscle circuit (Figure 2-4). We learn three things from these results. First, Hunchback controls circuit membership of motor neurons consistently across two lineages. Second, muscles that are over-innervated in comparison to control can have normal synaptic transmission, but muscles that are under-innervated in comparision to control have impaired synaptic physiology. Third, Hunchback controls circuit membership consistently across different time points in nerve cord development. This suggests that in many lineages Hunchback is likely to regulate circuit membership, which has broader implications extending to Hunchback’s vertebrate ortholog, Ikaros (Mattar et al., 2015) (Figure 8 A-C). Future work will need to determine the extent to which Hunchback’s function is conserved.

### Precocious expression of Castor or Pdm lead to permanent changes in motor neuron-to-muscle synaptic partnerships in Drosophila locomotor circuits

Hunchback is one member of a large class of molecules termed temporal transcription factors that in the Drosophila motor system also includes Castor (Cas) and Pdm. In vertebrates, a Castor homolog, Casz1, has also been shown to be a temporal transcription factor (Mattar et al., 2015). In this study, we asked to what extent can temporal transcription factors Cas and Pdm control circuit membership. Specifically, in NB7-1, precocious expression of the temporal transcription factor Castor (NB7-1>Cas) or Pdm (NB7-1>Pdm) leads to three different phenotypes: (1) Precocious expression of Cas in NB7-1 <u>*truncates*</u> the production of Eve(+) U motor neurons (Figure 5, Figure 8A-D). Notably, the U motor neurons that are produced are produced at the correct time in development. (2) In NB7-1>Pdm, in anterior segments A1-A3, there are <u>*late-born*</u> U5 motor neurons born at abnormally <u>*late times*</u> in development (Figure 6, Figure 8A-C). (3) In NB7-1>Pdm, in posterior segments A4-A7, there are <u>*late-born*</u> U4/U5 motor neurons born at abnormally <u>*early times*</u> in development (Figure 6, Figure 8A-D). For reference, previous manipulations that prolonged Hunchback expression in NB7-1 generated <u>*early-born*</u> U (U1) motor neurons at abnormally <u>*late times*</u> in development. Thus, using Cas and Pdm, we can probe the system to get a more comprehensive understanding of how changes in embryonic temporal identity, mediated by temporal transcription factors, impacts motor neuron-to-muscle synaptic partnerships.

We show that NB7-1 specific manipulation of both Cas and Pdm leads to permanent changes in motor neuron-to-muscle synaptic partnerships (Figure 5, Figure 6). These data show that, like Hunchback, Cas and Pdm act in the neuroblast as a molecular switch to determine how many neurons from the lineage will populate the dorsal muscle circuit. Therefore, our data support a model in which temporal transcription factors, as a class of molecules, control motor neuron circuit membership.

We note that manipulation of Pdm produces two different phenotypes (Figure 6). How does one manipulation give two opposing phenotypes? We find that the different phenotypes are correlated with A-P position, raising the possibility that this difference involves factors differentially expressed along A-P axis (e.g., Hox genes). For example, in anterior segments Pdm could work with one Hox gene to promote U4 molecular identity, whereas in more posterior segments Pdm could work with a different Hox gene to promote the same fate. Already, this type of differential combinatorial use of spatial and temporal factors has been shown to converge on a common neuronal molecular identity (Gabilondo et al., 2016). Notably, in the Gabilondo study, converging mechanisms occur in different neuroblasts lineages. Our data raise the possibility that converging mechanisms leading to one molecular identity could occur within one class of stem cell (e.g., NB7-1) found at different A-P positions.

In wild type, NB7-1 produces five U motor neurons, each of which can be uniquely identified by embryonic molecular marks, and each of which, in larvae, makes a unique motor neuron-to-muscle synaptic partnership. Notably, in NB7-1>Cas, the embryonic molecular identities that are lost are U4 and U5. In larval stages, the specific dorsal muscles that are under innervated are the normal U4 and U5 target muscles. In NB7-1>Pdm, in segments A1-A3, the U3 embryonic molecule identity is skipped, and in larval stages the normal U3 muscle is under-innervated. In NB7-1>Pdm, in segments A4-A7, extra U5 embryonic identities are produced, and in larval stages the normal U5 target is over-innervated. Thus, in both NB7-1>Cas and NB7-1>Pdm there is strong correlation between embryonic molecular identity and mature motor neuron-to-muscle synaptic partnerships. This finding is somewhat surprising given that in previous work, which prolonged Hunchback in NB7-1, there was a poor correlation between embryonic molecular identity markers and mature motor neuron-to-muscle synaptic partnerships (Meng et al., 2019). Notably the correlation between embryonic molecular identity and synaptic partnerships seen in this study using Cas and Pdm manipulation is more inline with the predominant model in the field. Specifically, the predominant model is that temporal transcription factors are master regulators of entire temporal programs, and should determine not just early neuronal features like marker gene expression, but also terminal neuronal features such as synaptic partner selection. Overall, our data demonstrate that there is variability in the predictive strength of molecular identity markers.

In this study, we generated “heterochronic” mismatches between a U motor neuron’s embryonic temporal identity and its birth time by precociously expressing Pdm in NB7-1. By studying heterochronic mismatches we can assess the relative contribution of intrinsic factors (e.g., neuronal gene expression) versus extrinsic factors (e.g., availability of synaptic partners or transient signaling cues) in control of synaptic partner choice. Our data in NB7-1>Pdm, support a model in which lineage intrinsic factors determine circuit wiring for neurons regardless of environmental cues present earlier or later within a small developmental time frame (Figure 8). This is in agreement with previous heterochronic mismatches that were induced by manipulation of Hunchback expression (Seroka and Doe, 2019; Meng et al., 2019). However, there are still two unanswered questions. First, the time scale of heterochronic mismatches studied so far is on the order of one to several neuroblast cell divisions, or hours. So it is unknown how heterochronic mismatches conducted at larger time scales would impact the system. Second, so far the heterochronic mismatches performed in the Drosophila motor system have largely focused on neuromuscular synapses as a readout. Neuromuscular synapses are distinct from neuron-neuron synapses in the CNS, in that they are usually one-to-one partnerships, and because motor neuron axonal targets are large pre-existing muscles. In the future, it will be interesting to understand the extent to which intrinsic versus extrinsic factors contribute to neuron-to-neuron synaptic partnerships.

## Conclusion

In conclusion, our data provide strong support for the hypothesis that temporal transcription factors as a class of molecules are highly potent regulators of circuit membership. Our data suggest that in the Drosophila motor system, motor neurons are born knowing which circuit they will populate and that this information is provided to them by temporal transcription factors acting in the neuroblast. It is possible that temporal transcription factors in other parts of the Drosophila CNS and vertebrate orthologs of the temporal transcription factors manipulated in this study regulate circuit membership decisions in similar ways. Together our work provides fundamental insight into the logic of motor circuit assembly, and has implications for evolution, medicine, and the genetic basis of behavior.

## Materials and Methods

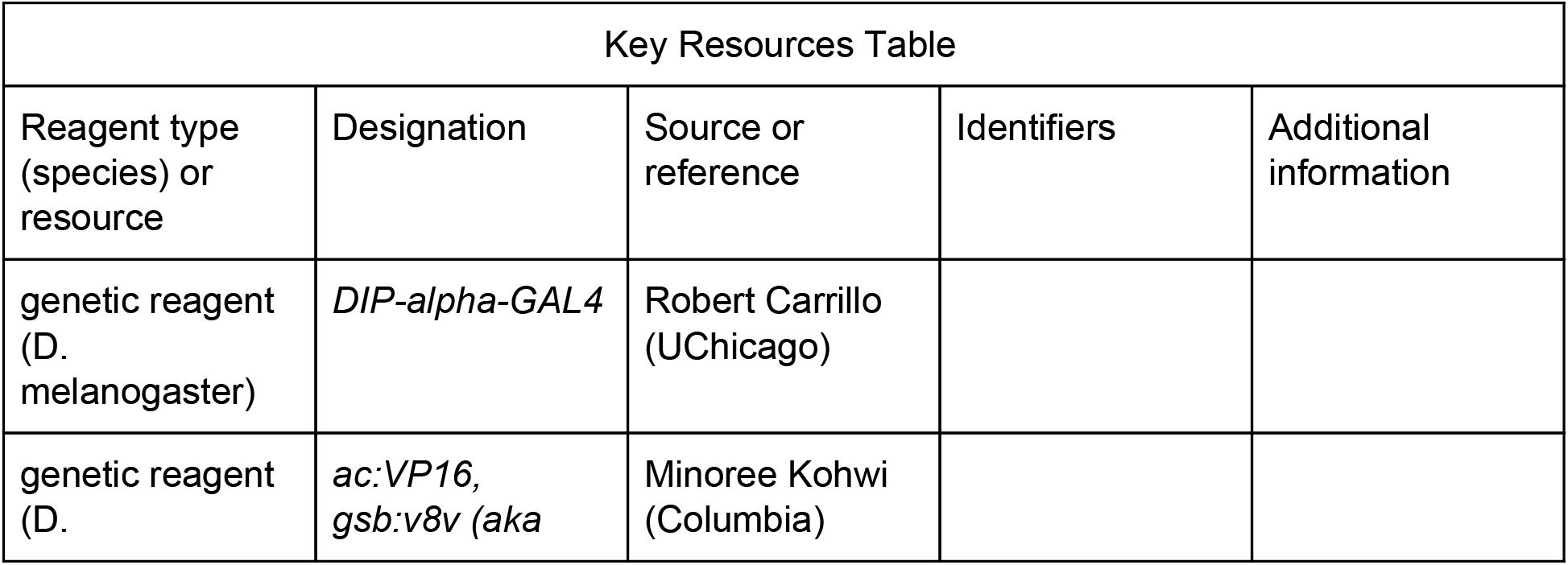
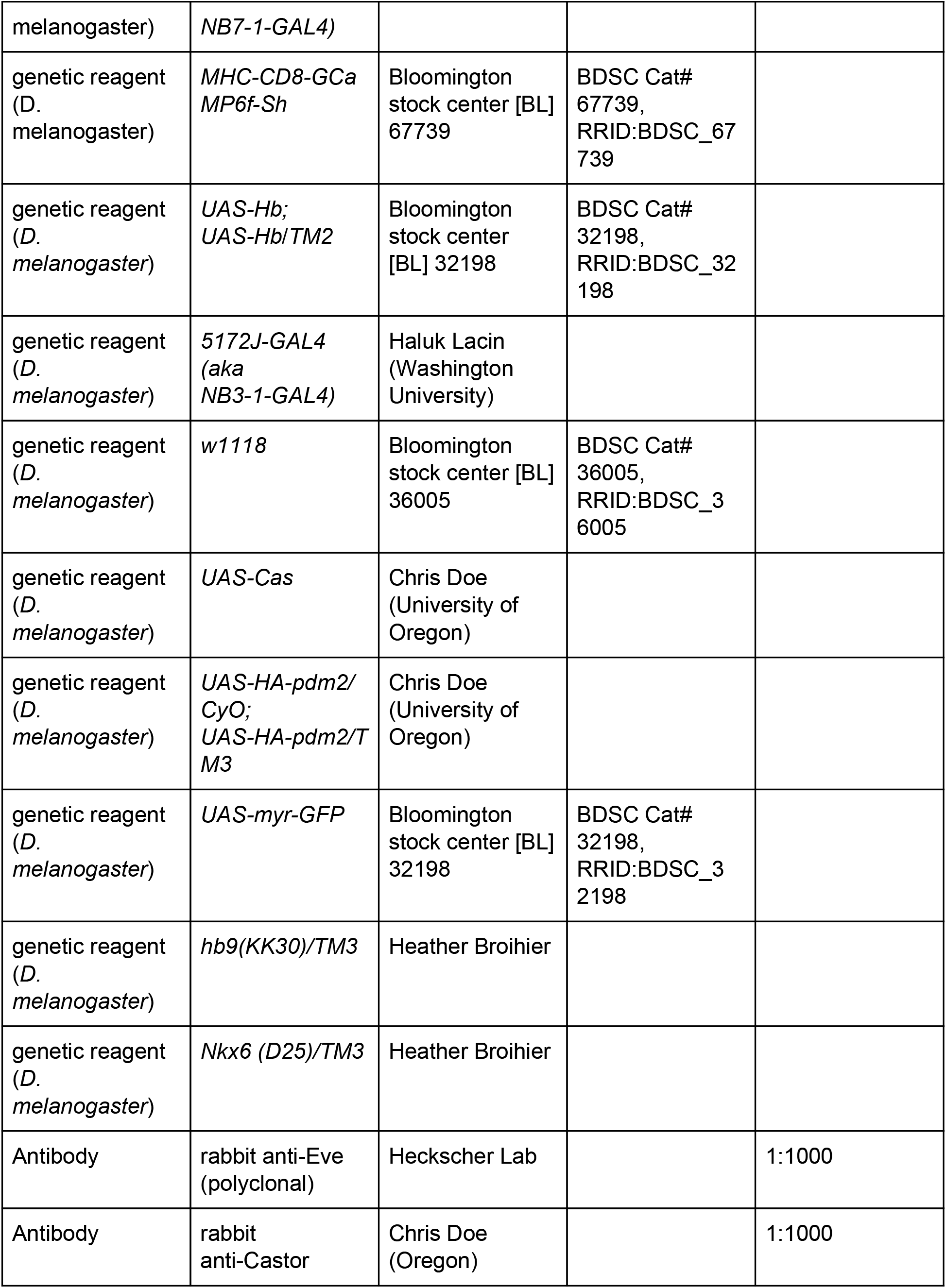
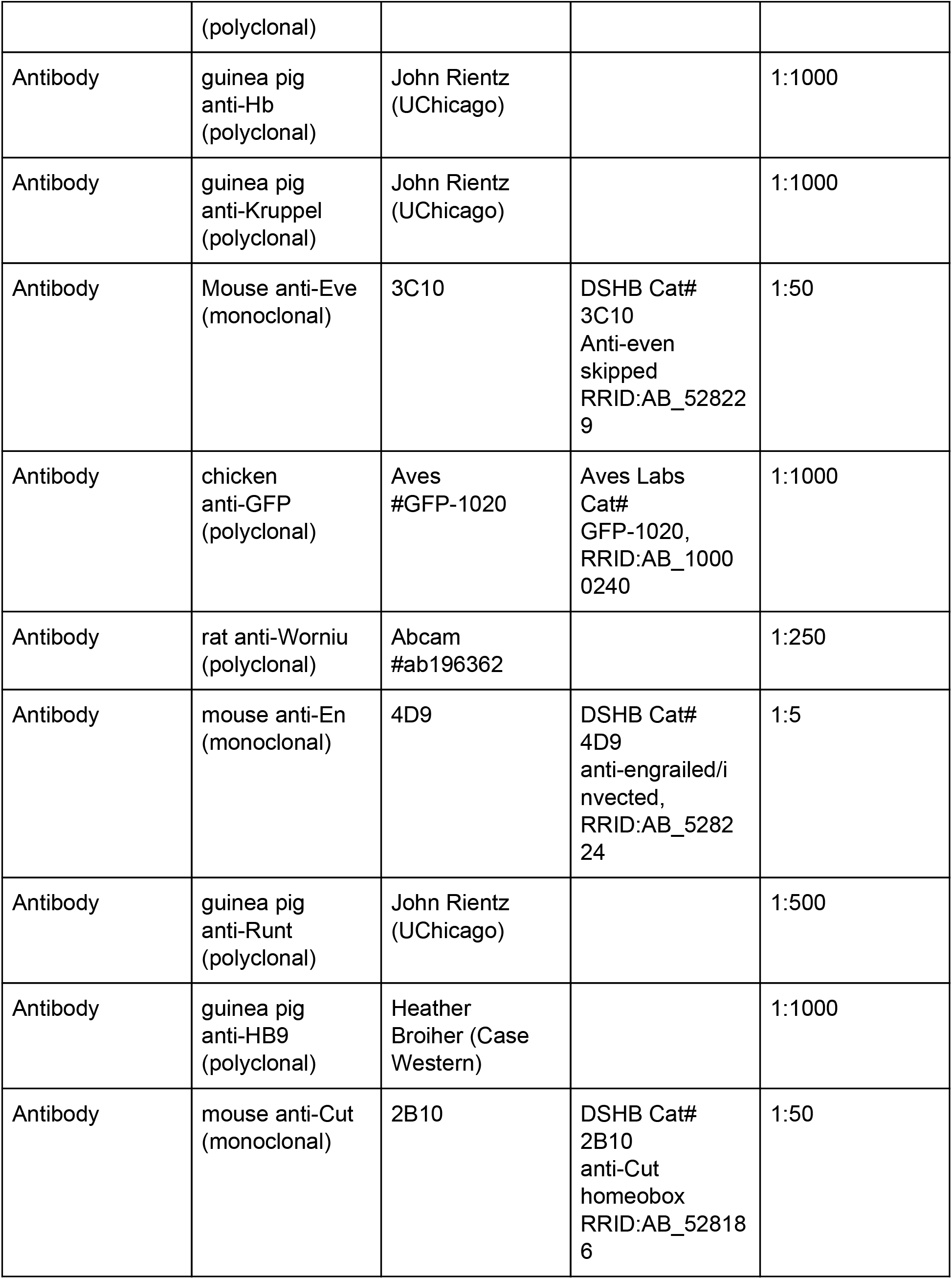
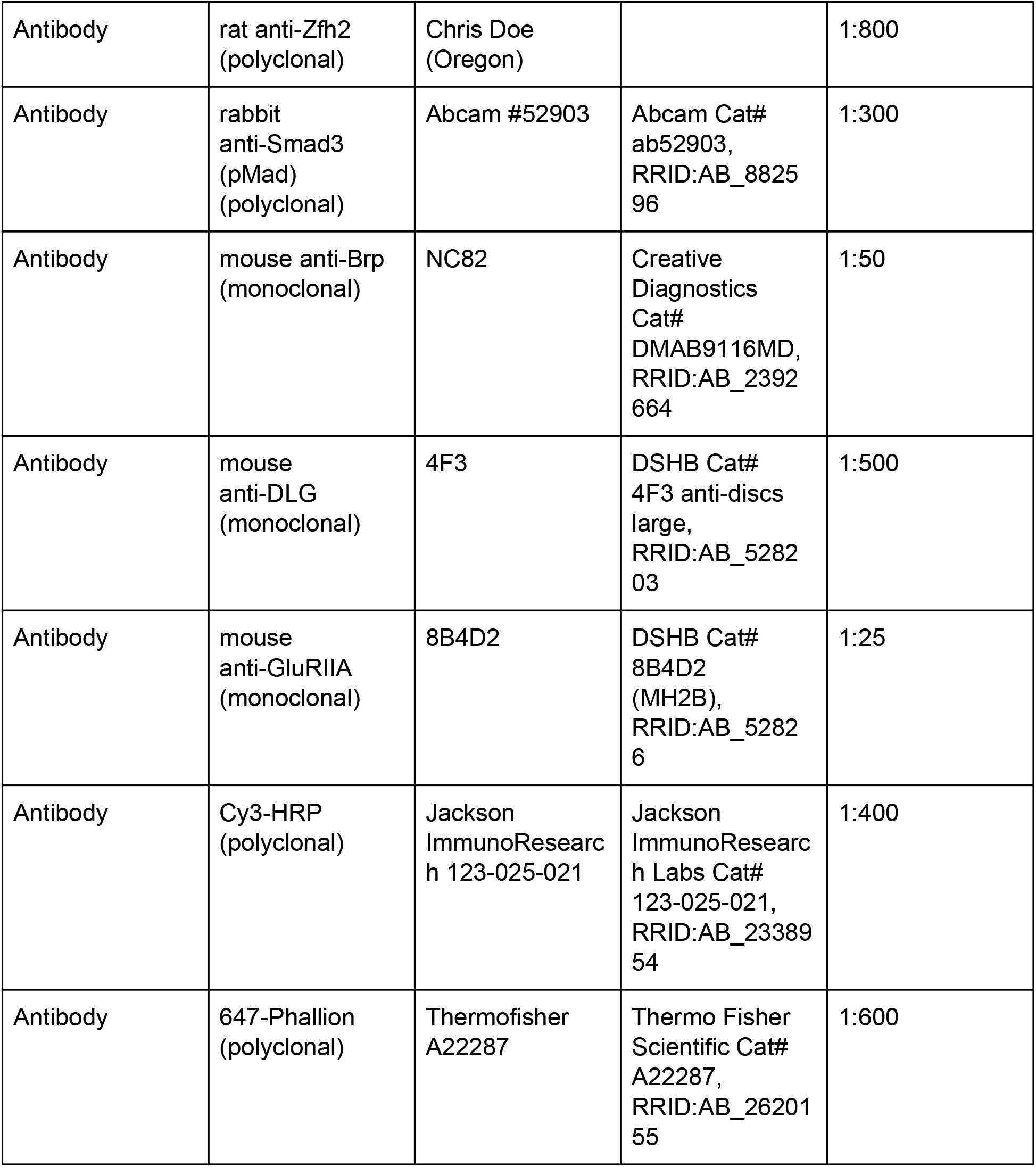

### Fly genetics

Standard methods were used for propagating fly stocks. For all experiments, embryos and larvae were raised at 25°C, unless otherwise noted. The following lines were used: *DIP-alpha-GAL4* (gift of R. Carrillo), *ac:VP16, gsb:v8v (aka NB7-1-GAL4)* (gift of M. Kohwi), *MHC-CD8-GCaMP6f-Sh* (Bloomington stock center [BL] 67739), *UAS-Hb; UAS-Hb*/*TM2* (BL 32198), *5172J-GAL4* (*aka NB3-1-GAL4*, gift of H. Lacin), *w1118* (BL 36005), *UAS-Cas* (gift of C. Doe), *UAS-HA-pdm2/CyO; UAS-HA-pdm2/TM3* (gift of C. Doe), *UAS-myr-GFP* (BL 32198), *Hb9(KK30)/TM3* (gift of H. Broihier), *nk6(D25)/TM3* (gift of H. Broihier).

### Tissue preparation

Three tissue preparations were used: Late stage whole mount embryos, in which antibody can still penetrate cuticle, and third instar (L3) fillet preparations, in which the neuromuscular tissue and cuticle are dissected away from other tissue and pinned open like a book, allowing for superb immuno-labeling and visualization of larval neuromuscular synapses. For all preparations, standard methods were used for fixation in fresh 3.7% formaldehyde (Sigma-Aldrich, St. Louis, MO) [49-51] or Bouin’s fixative (RICCA, Arlington,TX) for 7 minutes. For calcium imaging, L3 larvae expressing MHC-CD8-GCamp6f-Sh construct were dissected in HL3 solution containing 2mM Ca2+ and 25 mM Mg2+, brains removed, body walls rinsed with fresh saline, and samples imaged.

### Immunostaining

Tissue was blocked for an hour at room temperature or overnight at 4°C in phosphate buffered saline with 2% Normal Donkey Serum (Jackson ImmunoResearch), followed by 2 hours at room temperature in primary antibodies, and 1 hour at room temperature in secondary antibodies. Primary antibodies include: rabbit anti-Eve (1:1000, see Meng et al., 2019), chicken anti-GFP (1:1000, Aves #GFP-1020), rat anti-Worniu (1:250 Abcam #ab196362), guinea pig anti-Runt (1:500, John Rientz, UChicago), guinea pig anti-Hb (1:1000, John Rientz, UChicago), guinea pig anti-Kruppel (1:1000, John Rientz, UChicago), rat anti-Zfh2 (1:800 Chris Doe, UOregon), rabbit anti-Castor (1:1000 Chris Doe, UOregon), guinea pig anti-HB9 (1:1000 Heather Broiher, Case Western), rabbit anti-Smad3 (pMad) (1:300 Abcam #52903). The following monoclonal antibodies were obtained from the Developmental Studies Hybridoma Bank, created by the NICHD of the NIH and maintained at The University of Iowa, Department of Biology, Iowa City, IA: mouse anti-Brp (1:50, NC82), mouse anti-DLG (1:500, 4F3), mouse anti-GluRIIA (1:25, 8B4D2), mouse anti-En (1:5, 4D9), mouse anti-Eve (1:50, 3C10), mouse anti-Cut (1:50, 2B10). Secondary antibodies were from Jackson ImmunoResearch and were reconstituted according to manufacturer’s instructions and used at 1:400. 647-Phalloidin (1:600, Thermofisher A22287) Rhodamine-Phalloidin (1:600, Thermofisher R415), Cy3-HRP (Jackson ImmunoResearch 123-025-021), 647-HRP (Jackson ImmunoResearch 123-605-021). . Embryos were staged for imaging based on morphological criteria. Whole mount embryos and larval fillets were mounted in 90% Glycerol with 4% n-propyl gallate. Larvae brain preparations were mounted in DPX (Sigma-Aldrich, St. Louis, MO).

### Image acquisition

For fixed tissue images, data were acquired on a Zeiss 800 confocal microscope with 40X oil (NA 1.3) or 63 X oil (NA 1.4) objectives, or a Nikon C2+ confocal microscope with 40X (NA 1.25) or 60X (NA 1.49) objectives. For calcium imaging, data were acquired using a Zeiss 800 confocal microscope with a 40X dipping objective (NA 1.0) using 488-nm laser power with the pinhole entirely open. Images were acquired on a Zeiss 800 confocal microscope. Images were cropped in ImageJ (NIH) and assembled in Illustrator (Adobe).

### Image analysis

#### Type 1b branch counting

Third instar larval fillet preparations were stained for HRP to detect the neuronal membrane and Discs large (Dlg). Dlg allows type 1b and type 1s boutons to be distinguished. Single branches were counted as contiguous stretches of overlapping Dlg and HRP staining.

#### Calcium imaging

X-y-z-t stacks (Figure 9--figure supplement 1B-C) were converted into x-y time series images using the Maximum Intensity projection function (Fiji). X-y time series images were then registered using the Register Virtual Stack Slices plug-in (Fiji) to reduce movement artifacts. Time series images were projected into two different x-y single images using either the Maximum Intensity projection function (Fiji) or Average Intensity projection function (Fiji), and then the average intensity was subtracted from the maximum to get a change in fluorescence image.

### Electrophysiology

Third instar larvae were dissected and pinned in 0.3mM Calcium modified HL3 saline (Stewart et al., 1994)(70 mM NaCl, 5 mM KCl, 10 mM MgCl2, 10 mM NaHCO3, 5 mM Trehalose, 115 mM Sucrose, 5 mM HEPES). The ventral nerve cord was dissected away and the remaining body wall muscles were perfused with modified HL3 saline containing 0.5mM Calcium. In order to access muscle 30, muscles 6, 7 and 13 were gently removed using a sharpened tungsten wire. Preparations were visualized by a Nikon FS microscope using a 40×/0.80 water-dipping objective. Muscles in abdominal segments A3 and A4 were impaled by sharp electrodes (electrode resistance 15-30MΩ) filled with 3M KCl. mEPSPs were recorded, before stimulation was applied. The cut axon bundle was then stimulated by Master-9 stimulator (A.M.P.I.) at 0.2Hz to elicit EPSPs. Signals were processed by MultiClamp 700B (Molecular Devices), Digidata 1550B (Molecular Devices), and acquired by pClamp 10 software (Molecular Devices). Data was analyzed by Mini Analysis software (Synaptosoft). Recordings were rejected if the resting potential was > −60mV (for muscle 6) or > −50mV (for muscle 30). Quantal content was calculated by dividing the mean EPSP amplitude by the mean mEPSP amplitude from each muscle.

### Statistics

Descriptive statistics: average and standard deviation are reported, except for behavior, where the average of average speed and standard error of the mean are reported. Every data point is plotted in figures. Test statistics: All data was assumed to follow a Gaussian distribution. If standard deviations were unmatched Welch’s correction was applied. For numerical data in two populations, we used unpaired, two-tailed t tests. For numerical data in more than two populations, we used ordinary one-way ANOVAs with Dunnett or Games-Howell correction for multiple comparison. Analysis done using GraphPad Prism.

## Acknowledgements

We thank Chris Wreden, Marie Greaney, Zarion Marshall, Jake Henderson, Yi-wen Wang, Xiaoxi Zhuang, Johnathon Hall, Austin Seroka, and Chris Doe for comments on the manuscript. Minoree Kohwi, Haluk Lacin, Chris Doe, Heather Broihier, and John Rientz for fly stocks and/or antibodies. This work was funded by T32 GM007183 (JLM), NSF GRFP (DGE-1746045) (JLM), BSD International Student Fellowship (YW), NIH R01-NS105748 (ESH), University of Chicago MGCB start-up funds (ESH, RAC), National Institute of Neurological Disorders and Stroke K01 NS102342 (RAC).

## Competing interests

No competing interests declared

---

**Figure 2--figure supplement 1.**
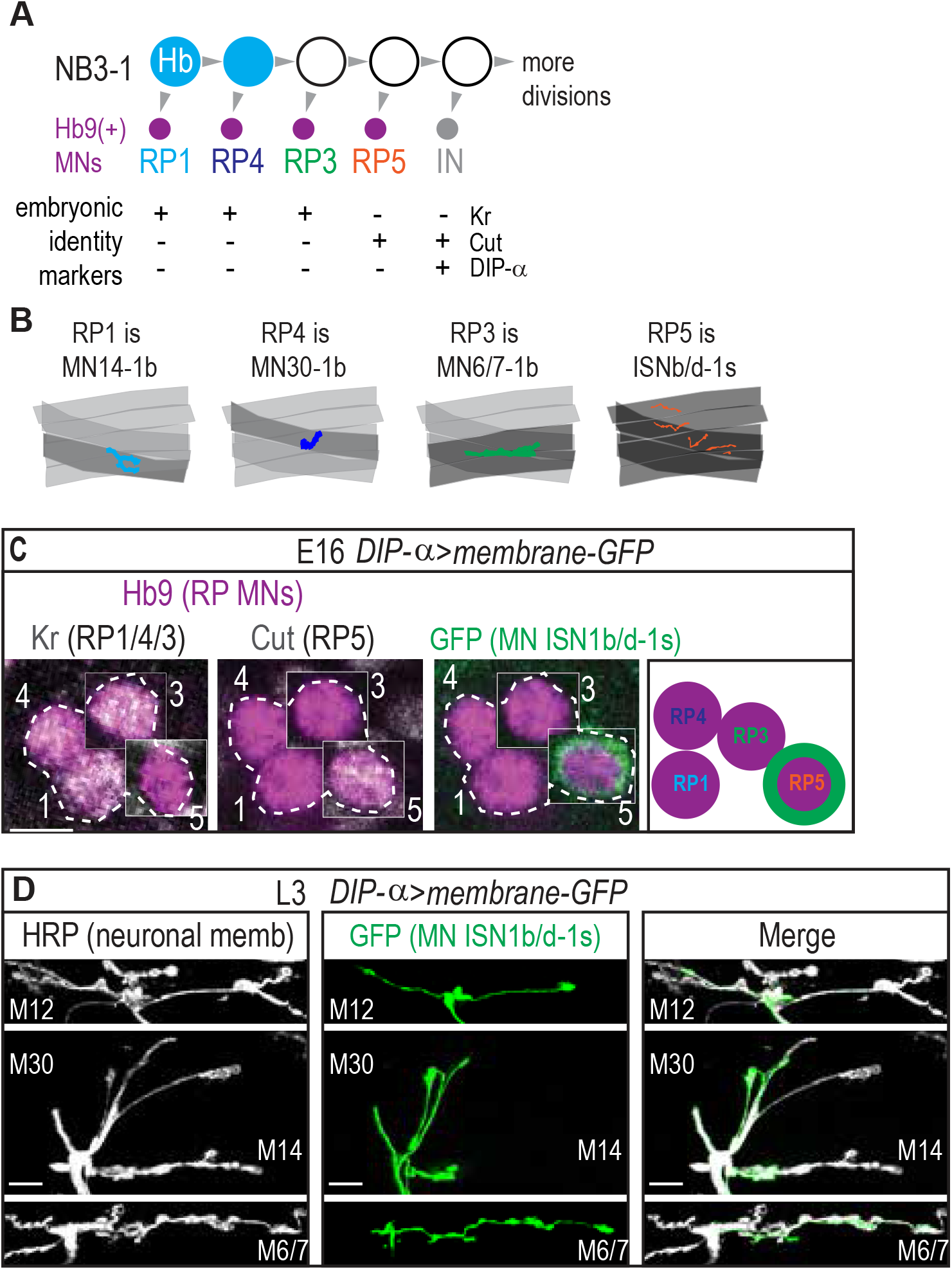
NB3-1 generates RP motor neurons, with known features. (A) Illustration of NB3-1 lineage progression. Each gray arrowhead represents cell division. Large circles are neuroblasts and smaller circles are neurons. Abbreviations: MN is motor neuron, IN is interneuron, Hb is Hunchback, and Kr is Kruppel. (B) Illustration of individual RP motor neuron neuromuscular synapses onto ventral muscles in third instar larvae. RPs populate the ventral muscle group motor circuit, which is independently recruited during locomotion. (C) Image of embryonic molecular identity marker and GFP expression driven by DIP alpha driver in Hb9(+) cell bodies in late stage embryonic CNSs. RP5 expresses GFP in late stage embryos. Boxes are insets from different z-planes (dotted outline around the RP motor neurons labeled by number). Scale bars represent 5 microns. Images are shown anterior up, midline to the left. (D) Images of individually labeled DIP alpha(+) motor neuron axon in third instar larvae (L3) fillet dissected larval prep. Scale bars represent 10 microns. Images shown in ventral view, anterior to the right, midline down. DIP-alpha>membrane GFP is DIP-alpha-GAL4/+; UAS-myr-GFP/+.

**Figure 2--figure supplement 2.**
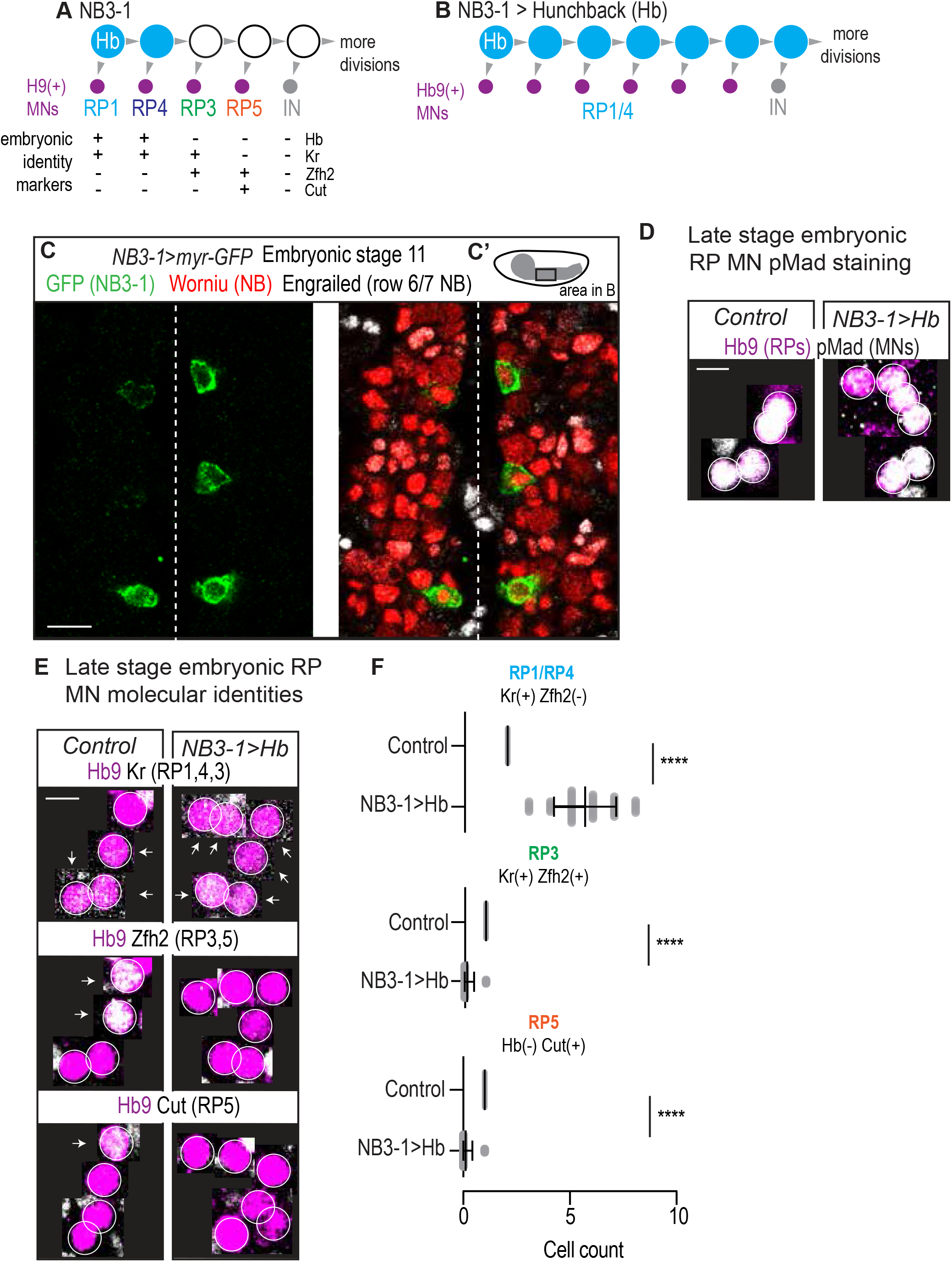
In embryos, prolonged expression of Hunchback generates more RP motor neurons with early born molecular identity. (A-B) Illustrations of NB3-1 lineage progression. Each gray arrowhead represents cell division. Each gray arrowhead represents cell division. Large circles are neuroblasts, and smaller circles are neurons. Abbreviations: IN is interneuron, Hb is Hunchback, Kr is Kruppel, Zfh2 is Zinc finger homeodomain 2. In NB3-1>Hb, there is an increase in the number of Hb9(+) with RP1/4 embryonic molecular identity. (C) Image of GFP in *NB3-1* as a reporter of NB3-1-GAL4 activity. Three complete abdominal segments are shown with anterior up, and midline dotted. C’ Illustration of a Drosophila late stage embryo, CNS in gray. For NB3-1>myr-GFP, n=28 hemisegments. Scale bar represents 10 microns. (D) Images of co-expression of Hb9 and the pan-motor neuron marker pMad in Control and NB3-1>Hb CNS of late stage embryos. For Control, n=160 hemisegments from 4 embryos. For NB3-1>Hb, n=360 from 6 embryos. Scale bar represents 5 microns. (E) Images of embryonic molecular identity marker expression in Hb9(+) cells in late stage embryonic CNSs. In NB3-1>Hb, extra Hb9(+) cells with RP1/4 molecular identity are produced. Boxes are neurons from different z-planes. Arrows indicate co-expression. Scale bar represents 5 microns. (F) Quantification of NB3-1 neurons in Control and NB3-1>Hb. Cells with RP1/RP4, RP3, and RP5 molecular markers. Each dot represents a single hemisegment. For Control, n=31,31,31 hemisegments from top to bottom graph. For NB3-1>Hb, n=60,90,60 hemisegments from top to bottom graph. NB3-1>myr-GFP is *NB3-1 GAL4/UAS-myr GFP.* Control is *W1118*. NB3-1>Hb is *NB3-1 GAL4/UAS Hb; UAS Hb/+*. Images are shown anterior up, midline to the left. For quantifications average and standard deviation are overlaid. ANOVA, corrected for multiple samples ‘****’ p<0.0001.

**Figure 5--figure supplement 1.**
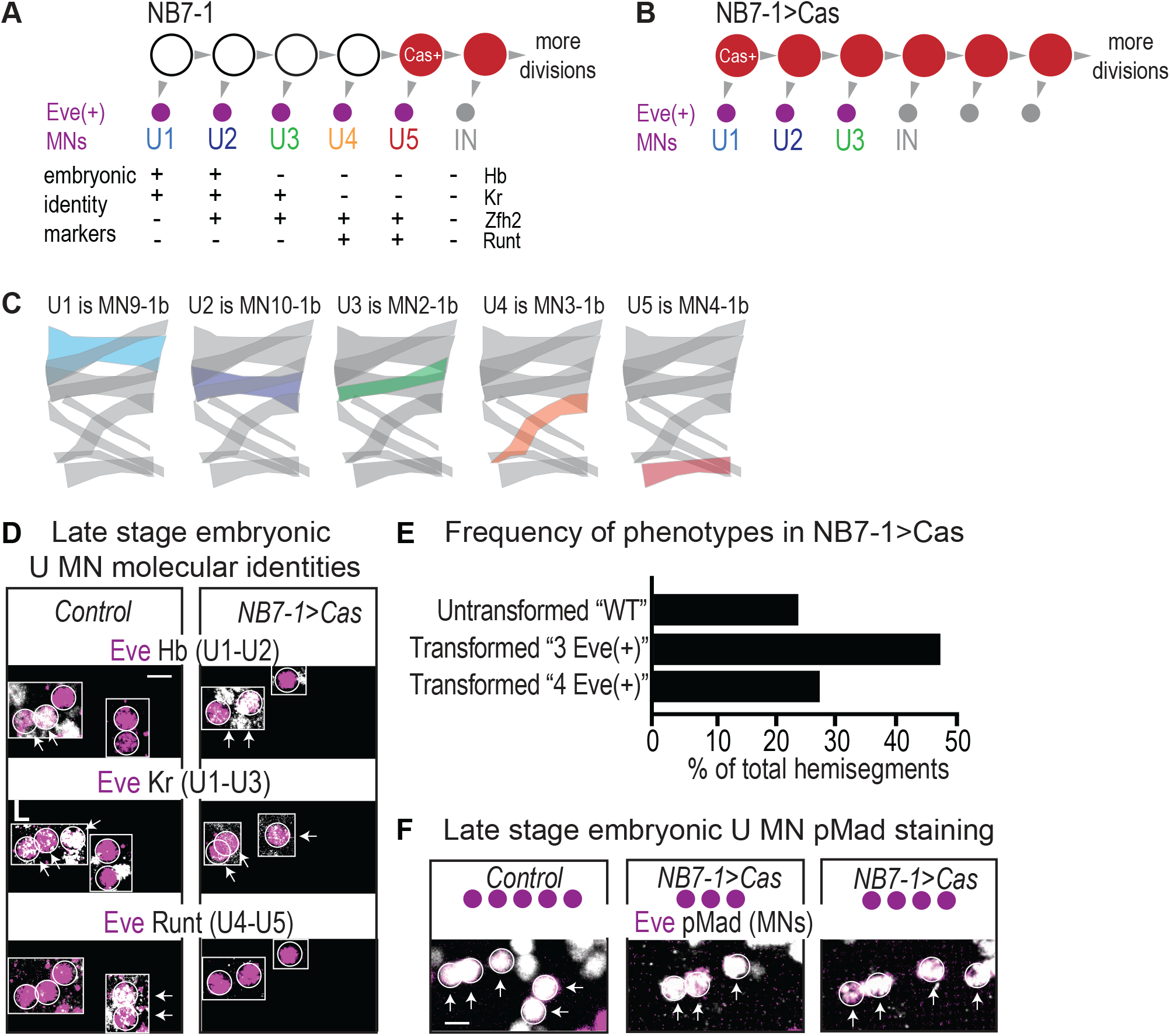
In embryos, precocious expression of Castor generates fewer U motor neurons at the expense of later born neurons. (A-B) Illustrations of NB7-1 lineage progression. Each gray arrowhead represents cell division. Each gray arrowhead represents cell division. Large circles are neuroblasts, and smaller circles are neurons. Abbreviations: IN is interneuron, MN is motor neuron, IN is interneuron, Eve is Even-skipped, Hb is Hunchback, Kr is Kruppel, and Zfh2 is Zinc finger homeodomain 2. In NB7-1>Cas, there is a decrease in the number of Eve(+) neurons with U4/U5 embryonic molecular identity. (C) Illustration of individual U motor neuron neuromuscular synapses onto dorsal muscles in larvae. Embryonic motor neuron (e.g., U1) and larval motor neuron synapse (e.g. MN9-1b) names are shown. Color code as in (A). (D) Images of embryonic molecular identity marker expression in Eve(+) cells in late stage embryonic CNSs. In NB7-1>Cas, Eve(+) cells with U4/U5 molecular identity are not produced. Boxes are neurons from different z-planes. Arrows indicate co-expression. (E) Quantification of the % of hemisegments in late stage embryonic NB7-1>Cas that give rise to five Eve(+) neurons (Untransformed “WT”), three Eve(+) neurons (Transformed “3 Eve(+)”), and four Eve(+) neurons (Transformed “4 Eve(+)”). n=44/180, n=86/180, n=50/180, respectively for the three phenotypes. (F) Images of co-expression of Eve and the pan-motor neuron marker, pMad in Control and NB7-1>Cas CNS of late stage embryos. All images are shown anterior up, midline to the left, scale bars represent 5 microns.

**Figure 6 --figure supplement 1.**
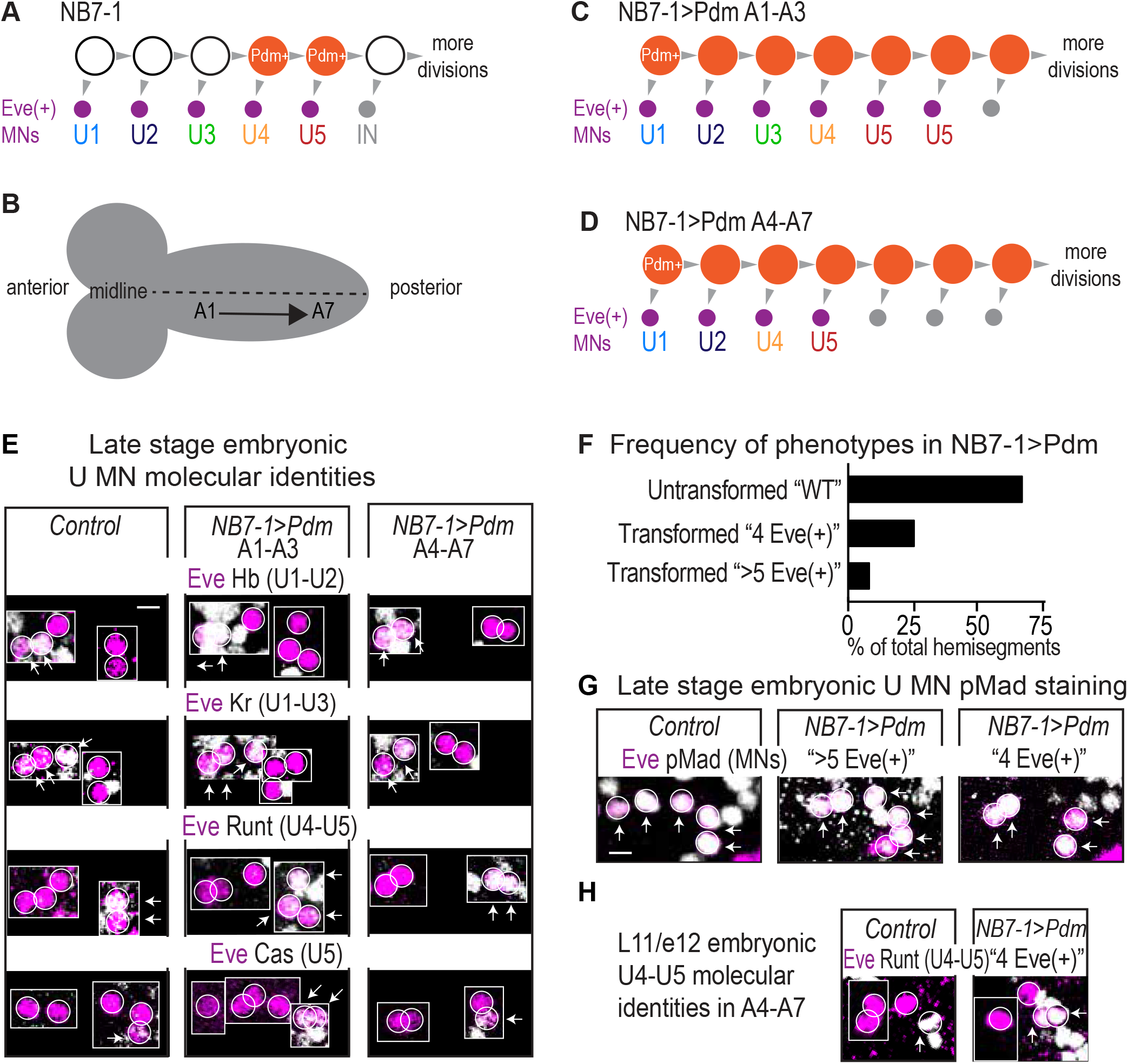
In embryos, precocious expression of Pdm generates either more or fewer U motor neurons depending on A/P positioning. (A) Illustration of NB7-1 lineage progression. Each gray arrowhead represents cell division. Each gray arrowhead represents cell division. Large circles are neuroblasts, and smaller circles are neurons. Abbreviations: IN is interneuron. (B) Illustration of a *Drosophila* embryo CNS. Nerve cord abdominal hemisegments are represented as (A). A1 is most anterior and A2-A7 are progressively more posterior. Dotted line represents the midline. (C-D) Illustration follows (A). In NB7-1>Pdm for A1-A3, there is an increase in the number of Eve(+) neurons with U5 embryonic molecular identity (C). In NB7-1>Pdm for A4-A7, there is a decrease in the number of Eve(+) neurons with U3 embryonic molecular identity. (E) Images of embryonic molecular identity marker expression in Eve(+) cells in late stage embryonic CNSs. In NB7-1>Pdm A1-A3, extra Eve(+) cells with U5 molecular identity are produced. In NB7-1>Pdm A4-A7, Eve(+) cells with U3 molecular identity are not produced. Boxes are neurons from different z-planes. Arrows indicate co-expression. (F) Quantification of the % of hemisegments in late stage embryonic NB7-1>Pdm that give rise to five Eve(+) neurons (Untransformed “WT”), four Eve(+) neurons (Transformed “4 Eve(+)”), and more than five Eve(+) neurons (Transformed “>5 Eve(+)”). n=103/154, n=39/154, n=12/154, respectively for the three phenotypes. (G) Images of co-expression of Eve and the pan-motor neuron marker, pMad in Control and NB7-1>Pdm CNS of late stage embryos. (H) Images of embryonic stage late 11 to early 12 (l11/e12) embryonic stage of co-expression of Eve and the U4/U5 marker Runt. In NB7-1>Pdm, cell division rate is not altered because four Eve(+) neurons are born, similar to Control. During L11 in NB7-1>Pdm, two neurons expressing Runt are born, while in Control only one neuron expressing Runt is born. Control is *Pdm/+* and NB7-1>Pdm is *NB7-1 GAL4/Pdm; Pdm/+*. All images are shown anterior up, midline to the left. Scale bars represent 5 microns.

**Figure 7--figure supplement 1.**
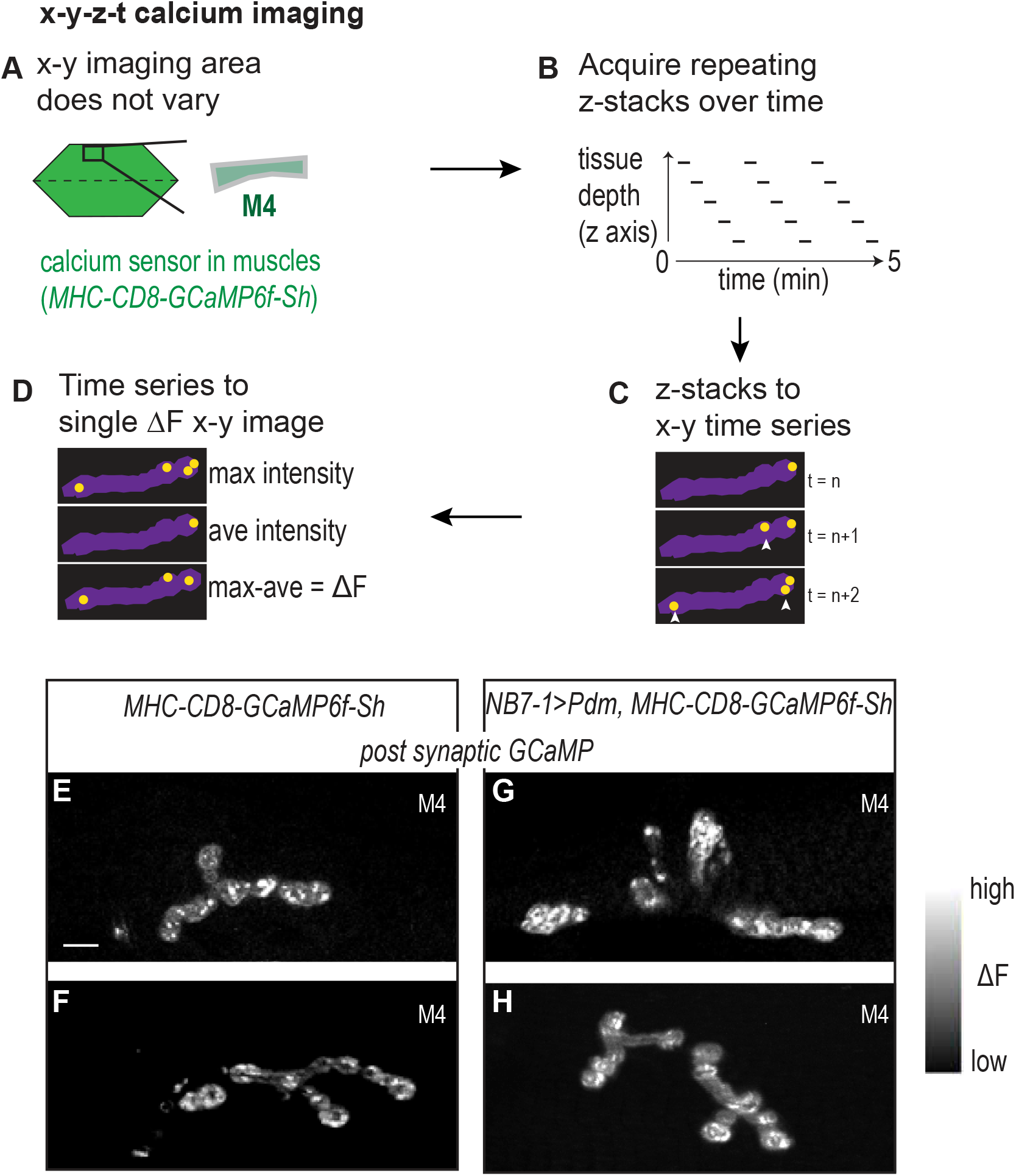
Calcium imaging protocol, analysis, and examples. (A-D) Illustration of calcium imaging protocol and analysis are shown. See Materials and methods for details. (E-H) Images of calcium signals on L3 Muscle 4. Control is *NB7-1-GAL4/+; MHC-CD8-GCaMP6f-Sh/+* and NB7−1>Pdm is *NB7-1-GAL4/UAS Pdm; MHC-CD8-GCaMP6f-Sh/UAS Pdm*. Images are dorsal up, anterior to the left. Scale bar represents 10 microns.

